# Transient centrosome loss in cultured prostate epithelial cells induces chromosomal instability to produce an oncogenic genotype that correlates with poor clinical outcomes

**DOI:** 10.1101/2025.05.10.653266

**Authors:** Jiawen Yang, Diogo de Oliveira Pessoa, John M. Ryniawec, Matthew R. Coope, Daniel W. Buster, Emily Loertscher, Mengdie Wang, Chen Chen, Anne E. Cress, Gregory C. Rogers, Megha Padi

**Affiliations:** Department of Molecular and Cellular Biology, University of Arizona, Tucson, AZ 85721, USA; Department of Cellular and Molecular Medicine, University of Arizona Cancer Center, University of Arizona, Tucson, AZ 85724, USA; Biostatistics and Bioinformatics Shared Resource, University of Arizona Cancer Center, University of Arizona, Tucson, AZ 85724, USA

**Keywords:** Chromosomal instability, centrosome, cancer evolution, prostate cancer, copy number variation, cancer signature, computational biology, multi-omics

## Abstract

Chromosomal instability (CIN) is a hallmark of prostate cancer that strongly correlates with metastatic burden and appears prominently in both primary cancer and metastatic disease. Low Gleason score primary prostate tumors display pervasive centrosome loss, a known mechanistic driver of CIN, that disrupts normal spindle assembly and increases mitotic errors. Previously, we found that transient depletion of centrosomes in immortalized, non-tumorigenic prostate epithelial cells (PrEC) induced a burst of CIN, generating cell lines capable of forming xenograft tumors. Here, we use a multi-omics approach to identify the oncogenic alterations caused by transient centrosome loss. By integrating genomic and transcriptomic data of the prostate lines, we identified a consensus set of focal copy-number variations (CNVs) induced by centrosome loss in cultured cells that are also detectable within a subset of samples from a prostate cancer patient cohort. Using this CNV signature, we were able to derive a unique transcriptomic signature from prostate cancer patient samples that showed strong predictive value for adverse clinical outcomes. In summary, our experimental system uses centrosome loss to promote a punctuated burst of genomic crisis that is characteristic of genome evolution during prostate cancer progression. Consequently, this prostate cancer model produced recurrent structural variations that are detectable in patient samples and associate with worse outcomes.

**Graphic Abstract:** 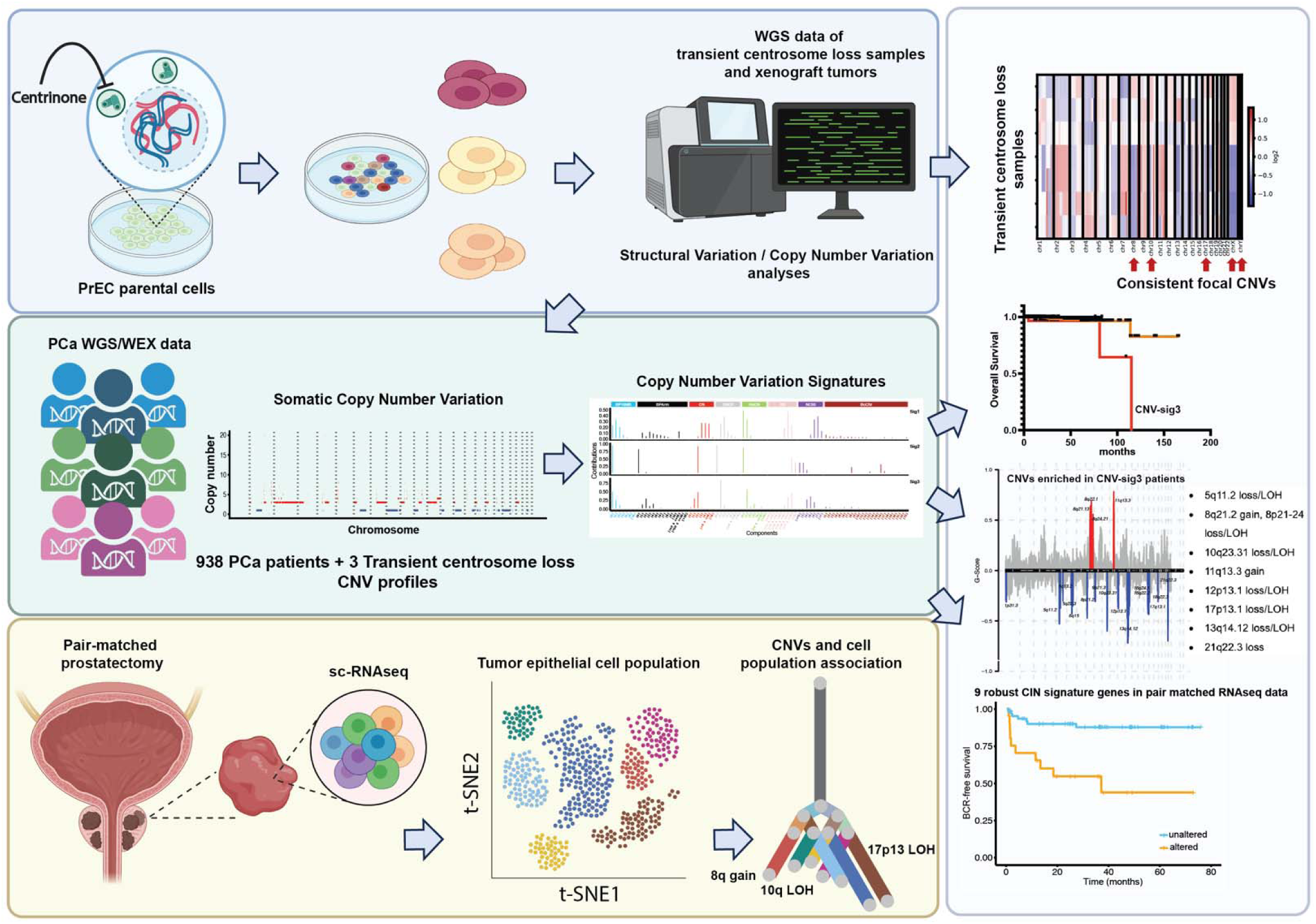

## INTRODUCTION

Prostate cancer (PCa) is the second leading cause of cancer death among American men and overtreatment is common due to the difficulty in distinguishing between indolent and potentially lethal cases. Unlike many cancers, PCa has a limited mutational burden (mean mutation frequency of 0.9 per megabase in the primary cancer genome) and common driver mutations are rarely detected in primary prostate tumors.^1–3^ Instead, localized, non-indolent PCa exhibits extensive chromosomal instability (CIN) with copy number variations (CNV), aneuploidy, and chromoplexy being the most common genetic changes.^2,4–9^ Indeed, CNVs and translocations are early events in prostate tumorigenesis.^10–15^ For example, both c-Myc and HER2/neu genes are amplified in precancerous prostatic intraepithelial neoplasia (PIN) ^16–19^, and approximately 15% of PIN lesions overexpress an oncogenic *TMPRSS2:ERG* gene fusion derived from an intrachromosomal translocation in chromosome 21.^10,20,21^ CIN is even more pronounced in metastatic PCa tumors; a recent survey of metastatic tumor genomes revealed that PCa has one of the strongest correlations between CIN and metastatic burden.^22^ These observations, plus the absence of identifiable driver mutations, suggest that CIN is the primary tumorigenic factor responsible for PCa.^2,23–25^ However, mechanistic drivers of CIN and genomic complexity in PCa are poorly understood.

Rather than a gradual accumulation of CIN during prostate cancer progression, genomic analysis indicates that tumor evolution occurs in a few successive bursts of CIN followed by expansion of the most adapted clones within the primary tumor, described as a model of ‘punctuated progression’.^5,9^ Punctuated progression is similarly observed early during triple-negative breast cancer progression.^26^ Furthermore, PCa evolution within the primary tumor leads to the emergence of aggressive sub-populations with metastatic potential.^27,28^ Indeed, metastatic populations can be tracked back to cells-of-origin within the primary tumor.^27,28^ It follows that prostate cells may experience short periods of genomic crisis that, consequently, drive tumor evolution. Therefore, we sought to recapitulate the punctuated burst model in a system where we could track the stepwise progression of these cells throughout their evolution to tumorigenesis. What then are the mechanisms that promote CIN in the prostate and potentially initiate bursts of genomic crisis?

Chromosome segregation errors during cell division are a primary cause of CIN.^29^ During mitosis, chromosomes are captured and segregated by the mitotic spindle. Importantly, the fidelity of chromosome segregation is governed by the shape of the mitotic spindle, so abnormally shaped spindles are error prone.^30^ Spindle shape is controlled by organelles known as centrosomes, microtubule-organizing centers that are positioned at both spindle poles.^31^ Normally, mitotic cells contain two centrosomes which guide assembly of a bipolar, fusiform-shaped spindle. However, changes in centrosome number (either too many or too few) result in abnormally shaped spindles that frequently missegregate chromosomes and promote different forms of CIN, including aneuploidy, polyploidy, translocations, and chromothripsis.^32–38^ The overproduction of centrosomes (a phenomenon known as ‘centrosome amplification’) is an emerging cancer hallmark.^39,40^

Previously, we found that cancer cells in low-grade (Gleason 3) regions of prostate tumors lacked centrosomes, a phenomenon that progressively worsened in higher-grade regions.^32^ Centrosome loss was subsequently described in epithelial ovarian cancers.^41^ Immortalized prostate epithelial cells experimentally depleted of centrosomes with centrinone, a reversible Polo-like kinase 4 (Plk4) chemical inhibitor,^36^ experienced mitotic errors that altered ploidy and increased the appearance of mitotic chromosome fragments, micronuclei, and multinucleated cells.^32^ Similarly, we detected multinucleated cells in low-grade primary tumors, 70% of which lacked centrosomes.^32^

We demonstrated that centrosome loss had the potential to transform non-tumorigenic, immortalized prostate epithelial (PrEC) cells, causing them to form xenograft tumors when injected into mice.^32^ To test this, PrEC cells were treated with centrinone for 15 days to transiently deplete centrosomes and induce a burst of genomic crisis, conceptually modeling the punctuated progression of PCa evolution. Single cells were then isolated for clonal expansion and centrinone was washed out. Importantly, centrinone washout allows centrosomes to reassemble, enabling the cells to maintain their altered genome (Figure 1A). Three clonal lines were then isolated, two of which formed large, solid tumors. While transient centrosome loss was sufficient to induce oncogenic transformation of PrEC cells, the differences between the clones’ tumorigenic capacity and inconsistent frequency of tumor formation suggested that centrosome loss induced unique molecular changes across cell lines. However, the genomic and transcriptomic alterations caused by centrosome loss-induced CIN in these lines are unknown.

**Figure 1.**
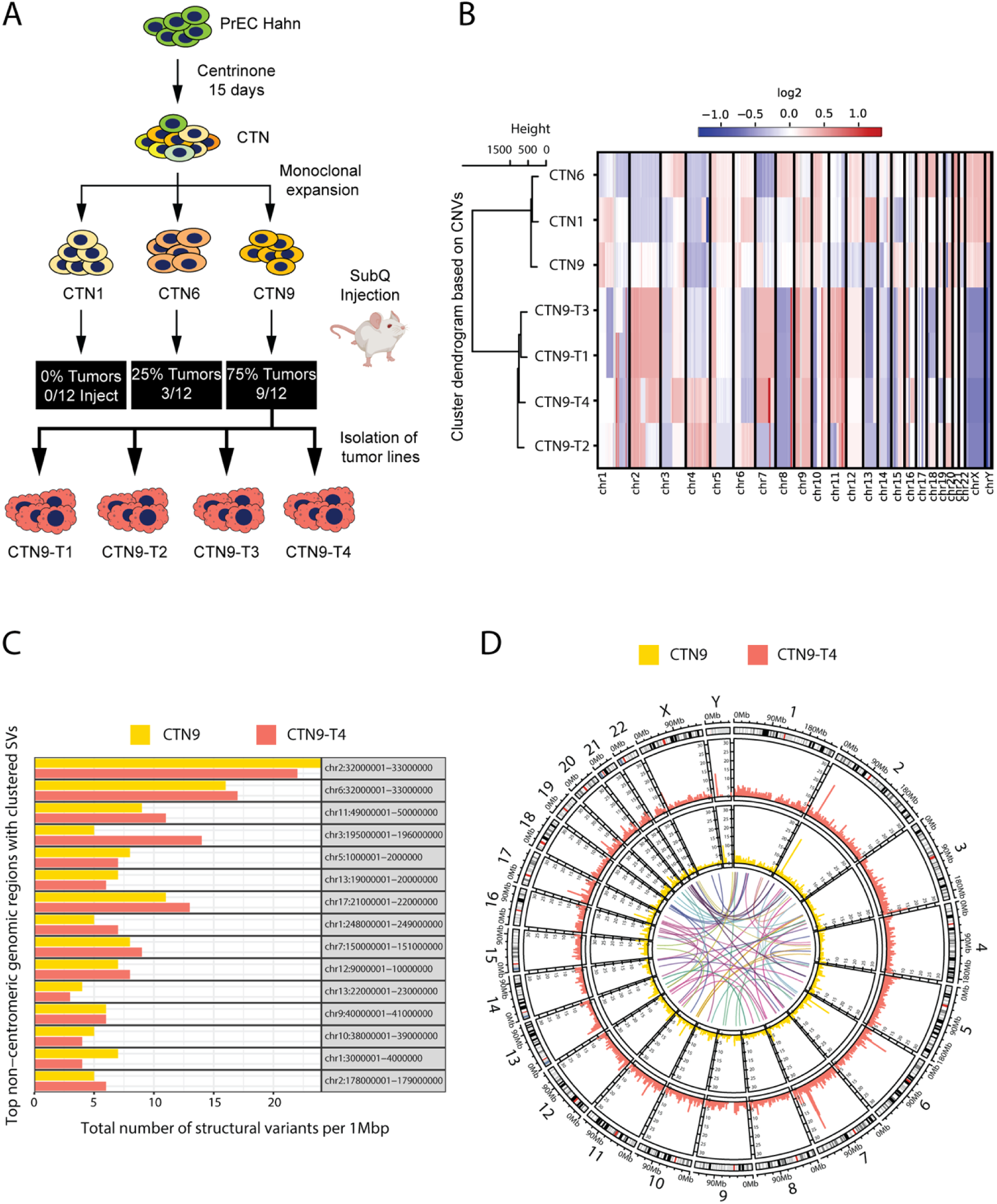
Genomic characterization of a transient centrosome-loss model of tumorigenesis. (A) Schematic of our transient centrosome loss model. The PrEC-Hahn line was treated with centrinone, a PLK4 inhibitor, for 15 days to block centrosome duplication and deplete centrosomes from the dividing culture. Following drug washout and centrosome reassembly, cells were clonally expanded and three lines (CTN1, CTN6, and CTN9) were tested in forming xenograft flank tumors in male mice. Tumor cells were isolated from four different CTN9 solid tumors (lines CTN9-T1, T2, T3 and T4). All cell lines were examined by whole genome sequencing and compared to parental PrEC. (B) Heatmap of somatic copy number variations (CNVs). Absolute copy number changes were transformed to log2 ratios [log2 (absolute copy number of a region/2)] with values above 0 indicating copy number gains (red) and values below 0 indicating copy number losses (blue). Samples were clustered based on Euclidean distance computed from region copy number data. (C) Bar plot showing non-centromeric SV hypermutated regions in CTN9 cells and one of its xenograft tumor lines, CTN9-T4. SVs were counted per 1 Mbp of the genome. (D) Circos plot displaying SVs across the entire genome for CTN9 (yellow) and CTN9-T4 (salmon). The center link plots depict both intra or inter chromosomal translocation events, while the inner and outer layer bar plots show the counts of SVs across every 1 Mbp of the genome.

Here, we use multi-omics analysis to elucidate the molecular alterations caused by transient centrosome loss that occurred during the tumorigenic process. We identified common focal CNVs that enriched within a tumorigenic subpopulation of cells in our model that are also present within patient samples. We then extracted a centrosome loss-associated CIN signature that shows strong predictive value at both molecular and clinical levels. Additionally, we found that recurrent focal CNV associated with transient centrosome loss are found in dominant clones in prostate tumors. Our results suggest that centrosome loss can initiate a punctuated burst of CIN to promote PCa progression and establishes a unique CIN signature that can stratify PCa patient outcomes.

## RESULTS

### A transient centrosome-loss xenograft model reveals a CIN-mediated tumorigenic process

Our previous study demonstrated that transient centrosome removal induced a burst of CIN sufficient to transform cells and produce xenograft tumors in mice, however, the specific genomic alterations underlying tumorigenesis were unknown (Figure 1A).^32^ In that approach, we treated immortalized, non-tumorigenic human prostate epithelial cells (PrEC) with centrinone to block centrosome duplication, effectively eliminating them from the population by dilution as cells divided over a 15-day period; by day 10, 90% of cells contain zero or one centrosome. Consequently, acentrosomal dividing cells experienced mitotic errors and CIN. Washout of the drug restored centrosome numbers, which, theoretically, would prevent further CIN and allow cells to then stably maintain their genomes. Ten clonal lines were isolated by single cell expansion and three lines were further tested for tumorigenicity (CTN1, CTN6, and CTN9) by subcutaneous injection into the flanks of NSG male mice. Notably, 25% of CTN6 injections and 75% of CTN9 injections generated solid tumors. Given its higher frequency of tumor formation, we sought to understand the molecular changes underlying the tumorigenic potential of the CTN9 line, therefore we isolated cell lines from four different CTN9 tumors: CTN9-T1, CTN9-T2, CTN9-T3, and CTN9-T4. Collectively, these lines were used to characterize the genomic alterations associated with transformation and tumor evolution. We performed whole genome sequencing (WGS) on the parental PrEC line, CTN1, CTN6, and CTN9, as well as the four CTN9-derived xenograft tumor lines. Somatic structural variations (SVs), which included copy number variations (CNVs), were determined by comparing all derived cell lines to parental PrEC to identify recurrent alterations using DELLY (SV caller) and CNVkit (CNV caller) methods.^42,43^

Genome-wide CNV log2 ratios demonstrated substantial genomic heterogeneity among CTN1, CTN6, and CTN9, including unique aneuploidy events within individual lines, which was expected given the apparent random nature of chromosome segregation errors caused by centrosome loss (Figure 1B). Notably, while CTN9 tumor lines (collectively referred to as CTN9-T) exhibited more CNV events than their parental CTN9 line (Figure S1A), they also showed increased sample purity, marked by a higher cellular prevalence of copy number changes compared to CTN9 (Table S1). This observation suggests that CTN9 experienced additional CIN during expansion, giving rise to a subpopulation of cells that were selectively enriched during xenograft growth. This evolutionary bottleneck however was absent when forming secondary tumors; CTN9-T xenografts grew much faster when reinjected into the flanks of male NSG mice (12 days until palpable tumor versus 65 days with CTN9) (Figure S1B). Moreover, CNV regions that emerged across all four CTN9-T lines with consistent gain or loss direction were defined as common focal CNVs. These common focal CNV regions indicate that they facilitated clonal sweeps of the population (Figure 1B). Common copy number gains were detected in regions such as 8q24.11-q24.21 (spanning *MYC*), 10q22.3 – q23.31 (spanning *TNKS2*, upstream of *KIF11*), 17q21.31, 17q25.3 (spanning *ITGB3*), and 21p11.2 (upstream of *USP25*) (Figure S1C; Supplemental Spreadsheet 1). Common copy number losses were observed in regions including 8p (spanning *NKX3-1*), 8q21-q23, 10p12 – p15, 10q11, 10q26.2-26.3, 17p11.1-p13.3 (spanning *TP53*), 17q12 (spanning *CDK12*), 17q21.2, 17q21.31-q21.33 (spanning *SPOP* and *NGFR*), 17q22 (spanning *TRIM37*), 17q24.1-q24.3 (spanning *CEP112*), 18q21.1 (spanning *MYO5B*), 18q22.31 (spanning *FBXO15*), 21q22.3 (*TMPRSS2-ERG* fusion associated), and Yq11-q12 (spanning multiple MSY genes) (Figure S1D; Supplemental Spreadsheet 1). Notably, several of these recurrent CNV regions have been reported in PCa datasets; 8p is among the most frequently deleted regions with 8p22 loss occurring in 29–50% of PIN, 32–69% of primary tumors, and 65–100% of metastatic cases ^44,45^. Additionally, CNVs associated with the loss of *TP53, SPOP*, and *CDK12*, as well as the gain of *MYC*, have been identified in large PCa cohorts ^2,23,46^. Thus, CTN9-T lines have limited interpopulation heterogeneity, appearing more similar to each other than the initial CTN9. The observed CNV patterns in the CTN9-T lines suggest that they were subjected to an evolutionary process in which CNVs conferring growth advantages were positively selected within the developing tumors.

Specific genomic regions, such as 17p11.2 (spanning *NCOR1*, downstream of *TP53*: 17p13.1), 13q11 (upstream of *BRCA2*: 13q13.1 and *RB1*: 13q14.2), and 12p13.32–p13.1 (spanning *CDKN1B* and *CHD4*) were identified as having the highest number of clustered SVs in CTN9 and that were passed to the tumor lines (Figures 1C and S2A, Supplemental Spreadsheet 2). Notably, these SV hypermutation regions associate with known PCa recurrent genomic altered loci (RGA) ^47^. In addition, we observed translocations of 21p11.2 and 21q22.11-q22.3 (spanning *ERG* and *TMPRSS2*) in CTN9 that were also inherited in the tumor lines (Figures 1D and S2B). Furthermore, high SV counts were located at or near centromeres on chromosomes 6, 10, 11, 13, 19-21 and Y (Figures 1D and S2B), a general phenomenon reported in the pan-cancer set that may impact kinetochore function (such as, spindle attachment or the Spindle Assembly Checkpoint response).^48^ Taken together, our findings demonstrate that transient centrosome loss induces CIN, leading to massive genomic alterations, including chromosomal gains, losses, and rearrangements, with the potential to oncogenically transform an immortalized epithelial line.

### CIN continues in proliferating CTN9 cells, producing a cancerous line

From our CNV data, we noticed several regions of loss and amplification that differed between CTN9 and the collective CTN9-T lines (Figure 1B). For example, copy number of the short arm of chromosome Y “Yp” was reduced in CTN9 compared to parental PrEC but completely lost in all four CTN9-T lines. Spectral karyotyping (SKY) of these cells confirmed that two copies of chromosome Y were present in most parental PrEC cells, however, most CTN9 cells contained one or zero Y chromosomes (Figures 2A, 2B, and S3A). In addition, bulk RNA-seq analysis confirmed loss of Y chromosome gene expression in the tumor lines compared to CTN9 (Figure 2C). Given these differences, we considered that CTN9 was genomically unstable and, after subsequent cell divisions, created new cell populations, including an oncogenic genomically-stable line that formed genomically similar xenograft mouse tumors (Figure 1B).

**Figure 2.**
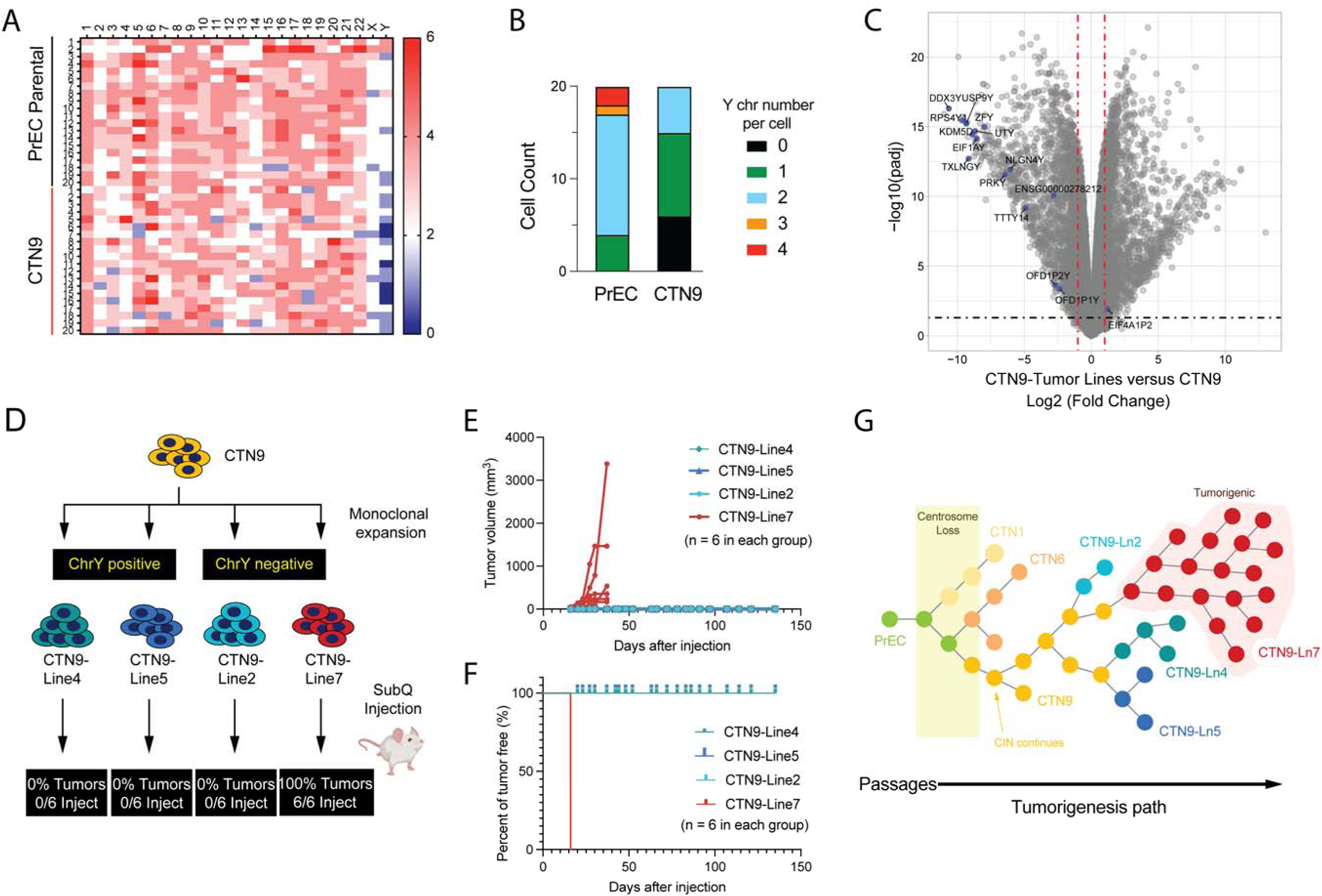
Genomic alterations in CTN-9 promote further CIN and heterogeneity, giving rise to an aggressive cancer type. (A) Heatmap of chromosome numbers derived from SKY analysis of PrEC parental and CTN9 cells. (B) Bar plot summarizing Y chromosome numbers in parental and CTN9 cells (*n* = 20 per condition). (C) Volcano plot comparing transcriptomic data between CTN9-tumor lines (T1, T2, and T3) and CTN9. Y chromosome-specific differentially-expressed genes (DEGs) are highlighted in blue. (D) Schematic shows second round of clonal expansion with CTN9, which contained aneuploid cells with loss of chromosome Y. Further xenograft tumor testing of four CTN9-derived lines identified only one line (#7) as positive. (E) Tumor volume growth curve for CTN9-Line2, 4, 5, and 7 following subcutaneous injection (*n* = 6 sites per line). (F) Kaplan-Meier curves showing tumor-free survival for each injection site of the four CTN9 derived cell lines. (G) Graphic summary of the transient centrosome loss model. PrEC cells experienced a brief period of genomic crisis due to a 15-day treatment of centrosome depletion, giving rise to new genotypes, including CTN1, 6, and 9. The genomic alterations in CTN9 increased the rate of CIN, producing further heterogeneity. Over time, CTN9 generates a new line (Ln7) which possesses oncogenic alterations. Most injections of CTN9 overcome the bottleneck to xenograft tumor formation, eventually producing genomically similar tumors due to population sweeps of CTN9-Ln7 in the cell population.

Therefore, we subjected CTN9 cells to a second round of single cell expansion and used chromosome Y status as a marker to potentially identify this oncogenic population. We isolated seven new clonal CTN9 lines (collectively referred to as CTN9-L) and then screened for loss of chromosome Y-specific genes using PCR (Figure S4A). We found that chromosome Y was retained in four of these lines but was lost in the remaining three. Fluorescence *in situ* hybridization (FISH) using chromosome Y-specific probes confirmed the PCR results (Figure S4B). We next selected four of these new clonal CTN9 lines for tumorigenicity testing: two lines that retained chromosome Y (lines CTN9-L4 and CTN9-L5) and two that lost chromosome Y (lines CTN9-L2 and CTN9-L7), resembling the CTN9 tumors. As before, cells were expanded and subcutaneous injected into the flanks of NSG male mice (Figure 2D). Strikingly, only CTN9-L7 formed tumors and did so at all injection sites (6 of 6) (Figures 2E and 2F). CTN9-L7 tumors were aggressive, appearing at 16 days (strikingly similar to CTN9-T lines) and eliminating the 65-day bottleneck displayed by its parent CTN9 line. Since CTN9-L2 did not form tumors, loss of chromosome Y in CTN9 was unlikely to be the transformative event. Furthermore, WGS of CTN9-L7 revealed that it exhibited a CNV pattern more similar to the CTN-T lines than CTN9, including copy number gain on 8q24.11- q24.21 and loss on 17p11.1 – p13.3, 17q12, 17q21.31-q21.33 and 17q22 (Figure S4D). These results confirm that CTN9 experienced continued genomic instability after transient centrosome loss to generate multiple subpopulations, at least one of which is oncogenic.

The genomic similarity of the CTN9-T lines and CTN9-L7 not only suggests that CTN9-L7 is the dominant clone that grew within the CTN9 xenograft tumors but that CTN9-L7 is more genomically stable than CTN9. Therefore, we tested if these cell lines display evidence of CIN to corroborate these predictions. First, we measured the frequency of micronuclei, which arise when mitotic errors generate lagging chromosomes during anaphase (Figure S3B).^49–51^ CTN1, CTN6, and CTN9 showed significantly elevated numbers of micronuclei compared to parental PrEC with CTN9 being the highest (Figure S3C). However, all of the CTN9 derivative cell lines (CTN9-L and CTN9-T) had significantly reduced micronuclei frequency compared to CTN9 (Figure S3D). To explain this increase in CIN, we measured centrosome numbers across the different lines because changes in centrosome number are known to promote chromosome missegregation. Cells were immunostained for both centriole (Cep135) and pericentriolar material (γ-Tubulin) components, as their co-localization marks bona fide centrosomes (Figures S3B). We found that centrosome loss was indeed a transient event as we designed it to be; most cells contained the proper two centrosomes in all of the lines (Figures S3E and S3F). Intriguingly, CTN9 had a significant subpopulation of cells with centrosome amplification (>2 centrosomes) and different from most of its derived lines, including CTN9-L7 (Figures S3E and S3F). Thus, our findings suggest that CTN9 has an elevated rate of CIN compared to parental PrEC and, although the underlying mechanism for this is unclear, CTN9 does possess a subpopulation of cells with centrosome amplification, a known CIN-generating mechanism that may be responsible for increased CIN.

Taken together, our findings suggest that transient centrosome removal induces a burst of CIN that can persist even after centrosomes are restored. However, genomically stable clones that acquire a selective advantage can emerge from this period of genome crisis and overtake the remaining population, recapitulating the punctuated features of PCa evolution (illustrated in Figure 2G).^5,11,31^

### A sequence of recurrent CNVs in PCa are common to CTN9-T evolution

We next wanted to test if the CNVs associated with progression of our model could contribute to tumorigenesis and tumor evolution in the prostate. Therefore, we analyzed an independent published single-cell RNA-seq dataset (GSE176031) from radical prostatectomy specimens that included pair-matched normal tissue.^52^ Data from four patients (PR5249, PR5251, PR5254 and PR5261) were integrated and annotated by cell origin (Figures 3A and 3B) and cell type (Figure 3C). Epithelial cells were selected to infer CNVs using the InferCNV algorithm (Figure 3D). Numerous epithelial populations in the PR5249 tumor sample exhibited high CNV amplitude in multiple regions including 3p25- p26 gain, 3q27-29 loss, 8q11.23 – q13.1, 8q24.13 – q24.3 gain, and 17p13.1-p13.3 loss (Supplemental Spreadsheet 3). Strikingly, several of the predicted CNVs in PR5249 aligned with the CNV regions identified in our transient centrosome loss model (Figure 3E), indicating a correlation may exist between these focal CNVs and PCa progression within this individual sample.

**Figure 3.**
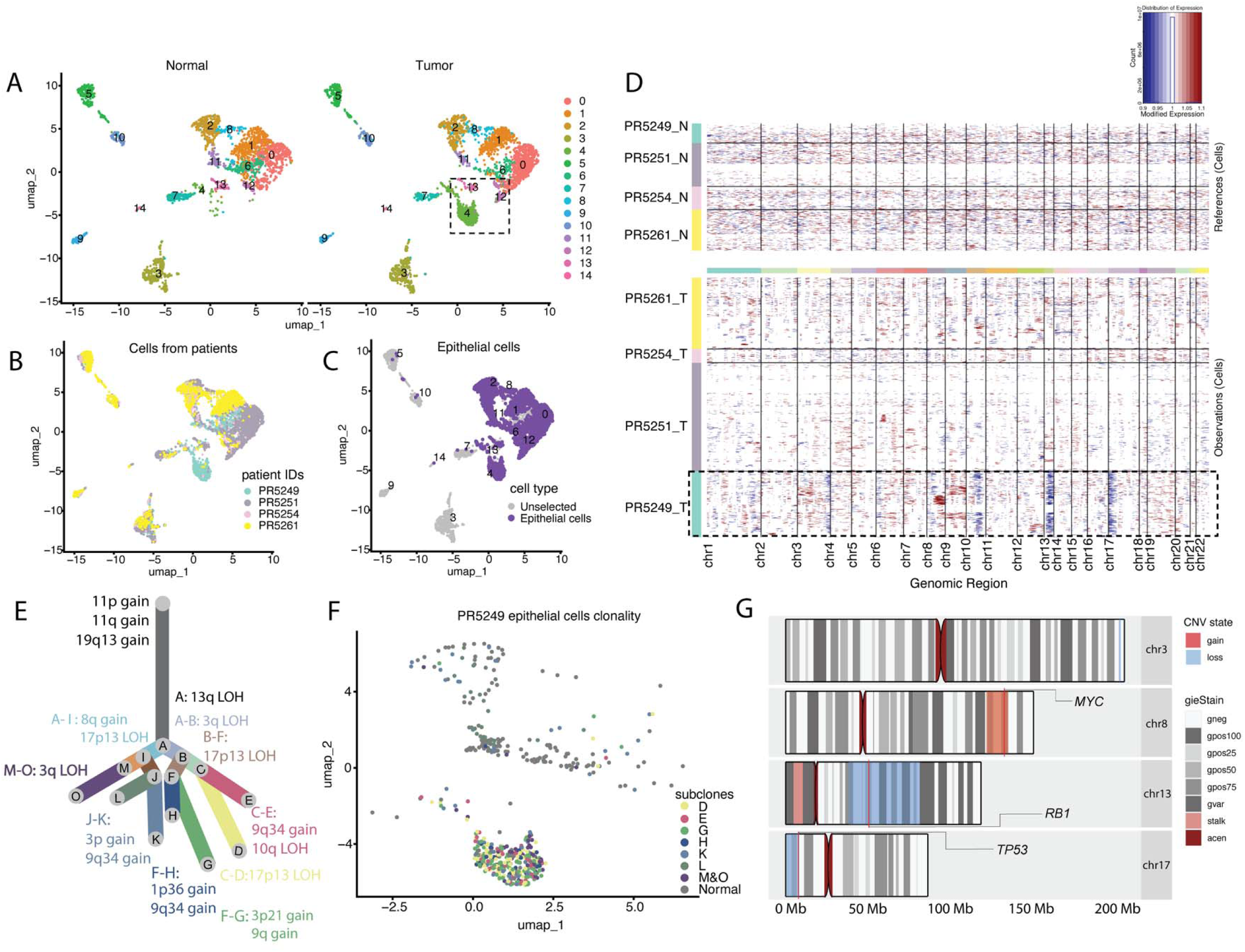
Single-cell analyses of PCa reveals focal CNVs similar to those in CTN9 tumor samples. (A) UMAP visualization of radical prostatectomy (RP) pair-matched samples from the GSE176031 dataset, annotated by sample origin (normal and tumor), and clustered based on gene expression profiles at a resolution of 0.4. The dashed square highlights cell populations unique to the tumor sample. (B and C) UMAP of the same dataset, annotated by (B) patient origin information and (C) their distinct gene expression profiles with epithelial cell populations highlighted in purple. (D) Heatmap of CNVs inferred from transcriptomic data of epithelial cells in RP tumor samples relative to epithelial cells in pair-matched normal tissue. Chromosomes X, Y, and mitochondrial DNA were excluded from the analysis. The dashed square highlights the PR5249 tumor sample which exhibits the highest genomic alteration among the four patient samples. (E) A phylogenetic plot constructed from PR5249 epithelial cells with branch lengths representing the population size of colonies. CNVs are color-coded to match the corresponding colony, illustrating the evolutionary relationship and genomic heterogeneity within the sample. (F) UMAP of PR5249 epithelial cells with colonies traced back from the phylogenetic tree in (E) and overlaid to visualize their distribution and relationship within the single-cell transcriptomic landscape. (G) Cytogenetic maps of chromosomes showing consensus CNV regions between transient centrosome-loss tumor samples and PR5249 tumor-associated epithelial cells. Amplifications are highlighted in orange and deletions are highlighted in blue. Consensus regions indicate shared CNV patterns across the CTN9 tumor samples.

We then applied Uphyloplot2 to the InferCNV results to understand the relationship between CNVs and cell population size within PR5249 (Figure 3F).^53^ Seven major colonies were detected from the PR5249 tumor epithelial cell population (only those with a population size ≥ 5% of total epithelial cell events were included) (Table S2). These populations co-segregated within the epithelial population (Figures 3A and 3F), and expressed high levels of PCa marker genes (*KLK2, KLK3,* and *ERG*), as well as low levels of epithelial cell markers (*KRT19* and *WFDC2*) (Figure S5A). Furthermore, these epithelial cell colonies expressed relatively high luminal cell markers, such as KRT8 and KRT18 and low basal markers such as ITGA6 and KRT5 (Figures S5B, S5C, S6A and S6B). These colonies also harbored genomic alterations associated with aggressive PCa phenotypes: colonies G and D contained alterations such as 17p13.3-p13.1 loss (*spanning TP53*), colony E contained 10q22.3 – q24.3 loss (spanning *PTEN*), colonies K and M-O contained 8q12.1 – q24.3 gain (spanning *MYC*), and all colonies contained 13q13.3 – q34 loss (spanning *RB1*) (Figure 3E, 3F and S6C). Furthermore, we identified progressive accumulation of focal CNV within the population, suggesting stepwise evolution of the genome from one colony to the next (i.e., colony D contains the CNV of colony C and 17p13 LOH) (Figure 3E). While potential branch points indicating divergence were identified, most of the cell population clustered within terminal colonies with few cells belonging to intermediary colonies (Figure 3E, Supplemental spreadsheet 3). Again, this may indicate a period of rapid genomic change followed by growth of advantageous, comparatively stable clones.

Several CNVs observed in the PR5249 tumor overlapped with those identified in our centrosome loss-induced xenograft tumors (CTN9-T) and were associated with PCa maker genes (Figures 3G and S7). Further examination of frequently reported PCa markers revealed that genes such as *ERG*, *PCA*3, *PTGR1*, and *AR* were highly expressed, whereas *PTEN* and *SPOP* showed low expression in the cell populations that co-localized with focal CNV-enriched colonies (Figure S6D). Taken together, our model of punctuated progression induced by centrosome elimination shares common genomic alterations with tumor evolution in a patient sample.

### A PCa-derived CNV-based CIN signature has predictive clinical value

We next asked whether the focal CNVs identified in our transient centrosome loss clonal lines bear any resemblance to CNV patterns found in larger PCa patient cohorts and potentially define a distinct genomic signature of CIN. Using CNV data generated from one of the CNV callers, Sequenza ^54,55^, we performed CNV signature analysis on 938 PCa patient samples ^2,3,23,56–58^, CTN1, CTN6, and CTN9 by employing Sigminer^55^, a method based on non-negative matrix factorization (NMF). To determine the optimal number of signatures, we employed the cophenetic correlation coefficient as a stability metric. We performed 50 independent NMF runs for each candidate signature number from *k* = 2 up to *k* = 12 to generate stable consensus clusters. After *k* = 3, we observed that the cophenetic correlation coefficient started decreasing rapidly, therefore three was chosen as the optimal signature number. The cumulative 941 samples were then divided into the optimal three distinct CNV signatures (Figures 4A and S8A). CTN1, CTN6, and CTN9 clustered in CNV signature 3 along with 236 patient samples (Figure 4A). Notably, CNV-signature 3 exhibited the highest proportion of metastatic tumor samples, with over 78% of the patients having metastatic PCa. In contrast, 62.6% of patients in CNV-signature 1 and only 5.62% of patients in CNV-signature 2 had metastatic PCa (Figure S8B). Given that CNV burden is positively correlated with tumor aggressiveness^4^, our CNV signatures likely reflects CNV burden to some extent; the CNV burden of signature 3 is significantly different from signature 1 and 2 (Figure S8C). However, the average CNV burden was similar in all three signatures (<1% of the genome) (Figure S8C), suggesting that CNV-signature 3 captures a distinctive CIN feature independent of burden.

**Figure 4.**
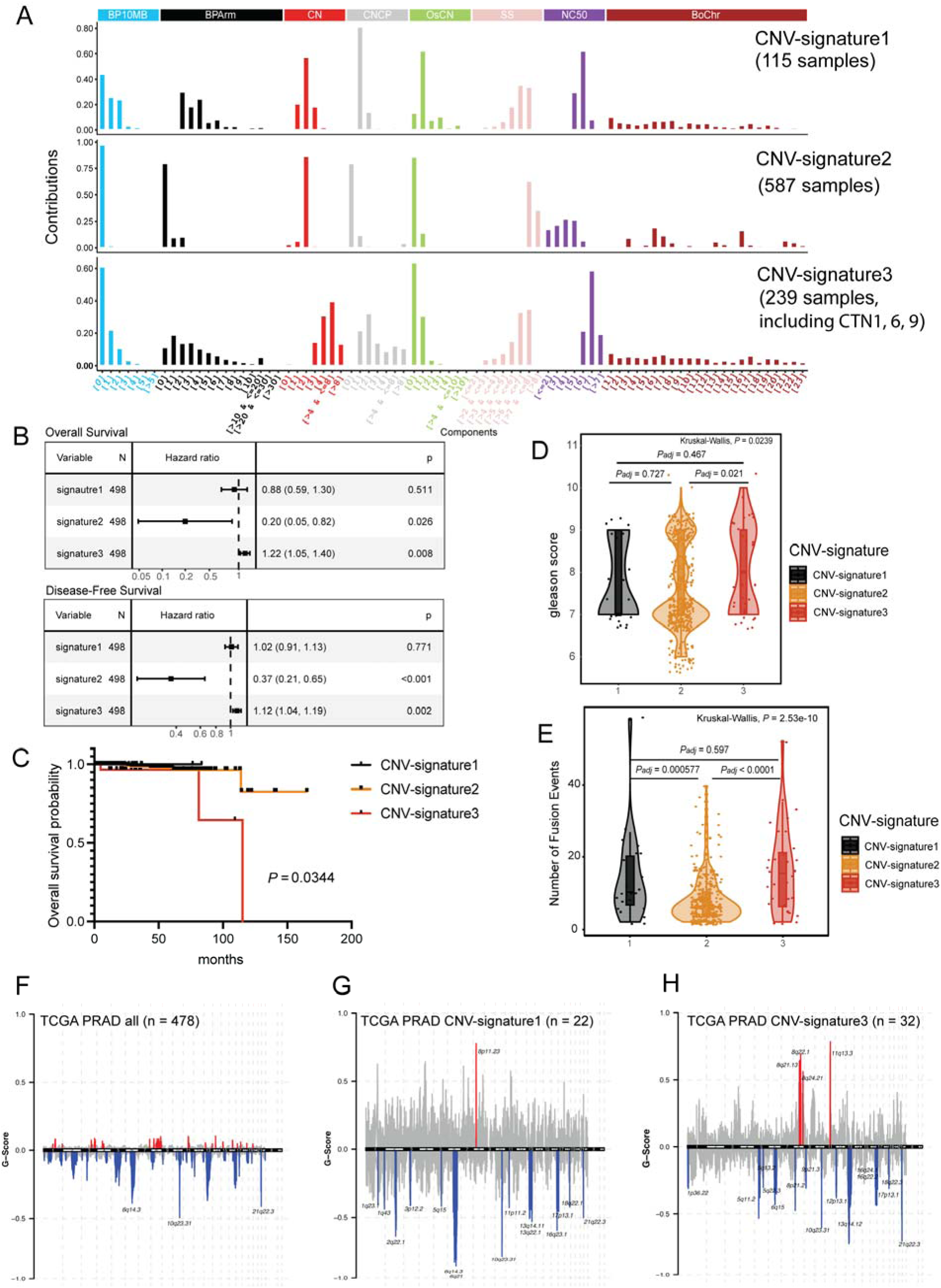
CNV signatures co-derived with transient centrosome loss samples have predictive clinical value in prostate adenocarcinoma (PRAD) (A) Three CNV signatures were identified from PRAD and transient centrosome loss cells. Each CNV signature comprises 8 distinct genomic features with a total of 80 components that were row normalized within each feature. (B) Forest plots show the relative risk of CNV signatures in 498 TCGA PRAD tumor samples for overall survival. Signatures were normalized to a range of 1-20 to evaluate the hazard ratio per 5% exposure increase. Hazard ratios and p-values were calculated using univariable Cox analysis. Squares represent hazard ratios and horizontal lines indicate the 95% confidence intervals. (C) Kaplan-Meier survival curves comparing overall survival for TCGA-PRAD samples attributed to each distinct CNV signatures. Statistical significance was determined using the log-rank test. The overall comparison showed significant differences among the groups (*P* = 0.0344). (D and E) Violin plots show (D) Gleason score (sum of primary and secondary site) and (E) number of fusion events in patient samples in the TCGA-PRAD dataset. Significance was analyzed using one-way ANOVA followed by Tukey’s HSD post-hoc test for pairwise comparisons with an overall difference observed among groups (D: F [2, 496] =3.763, *P* =0.0239; E: F [2, 428] = 23.28, *P* =2.53e-10). (F-H) GISTIC 2.0 results for TCGA-PRAD samples, including all samples (F) (*n* = 424), samples attributed to CNV-signature 1 (G) (*n* = 22), and to CNV-signature 3 (H) (*n* = 32). x-axis represents genome coordinates and y-axis shows G-scores calculated using GISTIC2.0 algorithm with the cut-off value of 0.9. CNV loss (G-score < 0) is shown in blue and gain (G-score > 0) shown in red.

CIN-associated PCa tumors typically exhibit an early-onset clonal chromosome complexity, driving rapid growth, metastasis, and early drug resistance.^59–62^ Identifying early diagnostic and stratification biomarkers to distinguish indolent from aggressive primary cancer is an important goal. To address this, we focused only on primary tumor samples and performed univariate Cox regression analysis to assess the impact of CNV-signature exposures on overall survival and disease-free survival in the TCGA PRAD cohort (Figure 4B). CNV-signature 3 significantly associated with worse survival outcomes, as indicated by a hazard ratio (HR) greater than 1 for both overall survival (HR = 1.22 [1.05, 1.40], *P* = 0.008) and progression-free survival (HR = 1.12 [1.04, 1.19], *P* = 0.002). In contrast, CNV-signature 2 was linked to improved survival with HR values significantly below 1 for both overall survival (HR = 0.20 [0.05, 0.82], *P* = 0.026) and progression-free survival (HR = 0.37 [0.21, 0.65], *P* < 0.001). CNV-signature 1 showed no significant association with either poor or improved survival outcomes. Kaplan-Meier survival analysis further confirmed that patients with CNV-signature 3 had significantly lower survival probabilities compared to other CNV-signatures (*P* =0.0034) (Figure 4C). Additionally, when comparing Gleason grade sums of primary and secondary lesions, patients with CNV-signature 3 displayed significantly higher-grade tumors than those with CNV-signature 2 (*P* = 0.0036) (Figure 4D). CNV-signature 3 was also associated with a higher number of gene fusion events compared to CNV-signature 2 (*P* = 9.7 x 10^-6^) (Figure 4E), further supporting its link to CIN. We obtained similar results when analyzing CNV data using an alternative CNV caller, FACETS ^63^ (Figure S9A-E; CTN samples were associated with signature 1 in these analyses).

To identify specific recurrent CNVs that may drive these signatures in TCGA PRAD patient samples, we performed GISTIC 2.0 ^64^ analysis across all 478 TCGA PRAD samples with available CNV information from the GDC data portal and were involved in signature analysis (Figures 4F-4H and S10A). GISTIC calculates a score for each CNV region considering both frequency in the population and CNV amplitude. Notably, regions such as 8q24.21 gain (spanning *MYC*), 11q13.3 gain (spanning *CCND1* and *PPP6R3*), 21q22.3 loss (*TMPRSS2:ERG* fusion-associated deletions), 17p13.1 loss (spanning *TP53*), 1p31.3 loss (spanning *JAK1*), and 13q14.2 loss (spanning *RB1*) were enriched in the TCGA PRAD samples associated with CNV-signature 3 compared to all samples (Figure 4H; Table S3). Regions including 8p11.23 gain, 2q22.1 loss, 6q21 loss (spanning *FOXO3*), and 10q23.31 loss (spanning *PTEN*) were enriched in the TCGA PRAD samples associated with CNV-signature 1 (Figure 4G). Importantly, multiple regions that were enriched in CNV-signature 3 samples overlapped with consensus CNV regions identified in our four CTN9 tumor lines (Figure S10B; Table S4). Lastly, we examined the relationship between the CNV-signature 3 and another indicator of genomic instability, the Microsatellite Instability (MSI) score. The MSI scores were previously computed using the MANTIS algorithm.^65^ Our analyses demonstrated that samples classified under CNV-signature 3 exhibited significantly higher MSI levels compared to those in CNV-signatures 1 and 2 (*P* = 0.02557). Moreover, CNV-signature 3 exposure scores showed a positive correlation with MSI scores (r = 0.3, *P* = 0.024), whereas CNV-signature 2 exposure scores were negatively correlated with MSI scores (r = −0.32, *P* = 0.015) (Figure S11A-D).

### Gene expression features associated with CNV-signature 3 predicts poor outcomes in independent PCa cohorts

We next sought to identify a corresponding transcriptomic signature that reflects the molecular changes associated with tumors in CNV-signature 3 using paired RNA sequencing data. Using LassoCV with 10-fold cross-validation to optimize the regularization parameter (λ) on TCGA PRAD bulk RNA-seq data, we identified a robust set of nine genes (*PRR11*, *PPP6R3*, *ACOT2*, *SHC2*, *SMC2*, *KIF11*, *POLE3*, *TMEM183A*, and *ZNF667* [Table S5]) that best distinguished CNV-signature 3 samples from CNV-signatures 1 and 2. Some of these genes have well-established functions in maintaining genomic stability and are mis-regulated in cancer, such as *KIF11* (mitotic kinesin-5 motor protein), *SMC2* (condensin I and II subunit), and *POLE3* (DNA polymerase epsilon 3).^66–71^ We then tested the prognostic value of this gene signature in three independent PCa patient transcriptomic datasets from German Cancer Research Center (DFKZ), Stand Up to Cancer (SU2C/PCF), and Memorial Sloan Kettering (MSK) by stratifying samples into two groups based on whether their gene expression z-score was extremal (|z|>2 or |z|>3) ^47,72,73^. To ensure reasonable sample sizes across datasets, we applied dataset-specific thresholds that accounted for the distribution of expression levels of the nine genes (|z|>2 for DKFZ and SU2C/PCF datasets, |z|>3 for MSK dataset).^23,47,72^ Our analysis revealed that dysregulation of the nine CIN signature genes (CIN9, hereafter) was significantly associated with poor clinical outcomes within three different PCa patient studies, which included reduced biochemistry recurrence (BCR)-free survival, disease-free survival, and overall survival (DKFZ: BCR-free survival, *P* = 0.00012, MSK: Disease-free survival, *P* = 0.0068, SU2C/PCF DREAM TEAM: overall survival, *P* = 0.046) (Figures 5A-C). Additionally, tumors with dysregulation of the CIN9 signature genes generally exhibited higher Gleason grades (Figures 5D and 5E). Within individual datasets, dysregulation of CIN9 genes were also associated with (i) a higher tumor purity (Figure 5F), which is a characteristic of early-onset tumors driven by CIN, (ii) a higher proportion of aggressive tumor types (Figure 5G), and (iii) a greater fraction of altered genome (Figure 5H). Lastly, we compared the performance of our CIN9 gene signature to five established CIN gene signatures: CIN25, CIN70, CIN23, CIN7, and CA20.^74–77^ We found that CIN9 exhibited more consistent predictive performances among all these CIN gene signatures when tested in the DKFZ, MSK, and SU2C/PCF datasets (Figures 5A-5C and S12). Strikingly, CIN9 was the only signature that maintained robust predictive power in the SU2C/PCF PCa cohorts (Table 1). Thus, our transcriptional CIN signature (CIN9), derived from pair matched genomic data in PCa samples that contain CIN, outperforms the predictive power of many other established CIN signatures.

**Figure 5.**
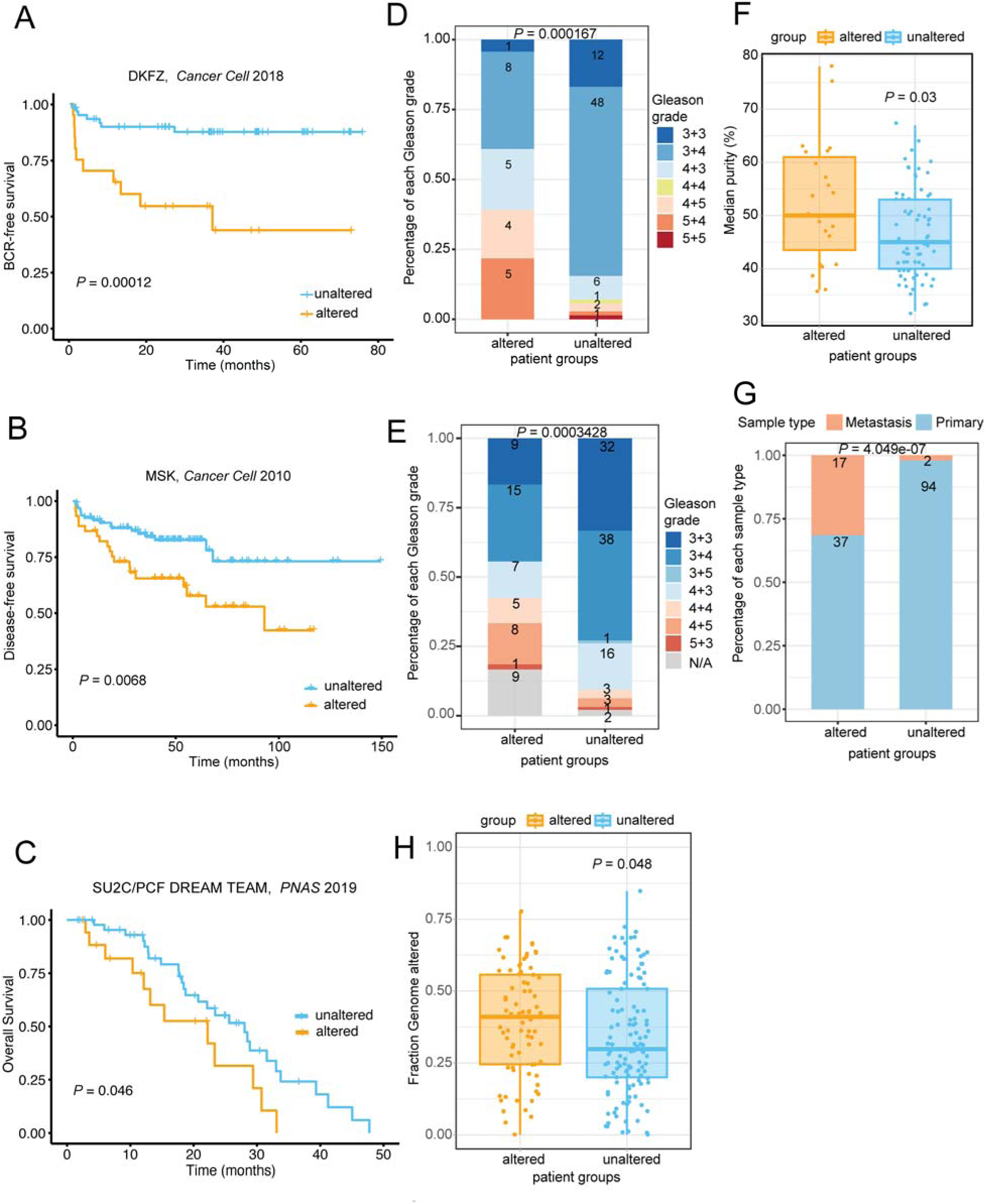
A CNV-based transcriptional CIN signature predicts poor outcomes in independent PCa cohorts. (A-C) Kaplan-Meier survival curves comparing (A) biochemical recurrence-free survival between signature gene-altered and unaltered patient groups in the DKFZ PCa cohort (*n* = 20 altered, 61 unaltered)^47^ (B) disease-free survival in the MSK PCa cohort (*n* = 23 altered, 71 unaltered)^72^, and (C) overall survival in the SU2C/PCF DREAM TEAM PCa cohort (*n* = 18 altered, 47 unaltered)^83^. Log-rank test was used to compare groups. (D and E) Bar plots show Gleason grade distribution based on signature genes in (D) *n* = 23 altered and *n* = 71 unaltered groups in DKFZ PCa cohort, and (E) *n* = 54 altered and *n* = 96 unaltered groups in MSK PCa cohort. Significance calculated with Fisher’s exact test. (F) Bar and whisker plot shows tumor sample purity in signature genes altered (*n* =23) and unaltered (*n* = 71) groups in DKFZ PCa cohort. Significance determined using student’s t-test. (G) Graph shows tumor sample types (primary and metastatic tumors) in signature genes altered (*n* = 54) and unaltered (*n* = 96) groups in MSK PCa cohort. Significance determined using Fisher’s exact test for the altered group with higher proportion of metastatic samples. (H) Bar and whisker plot shows percent genome altered (PGA) in signature genes altered (*n* = 90) and unaltered (*n* = 111) sample groups in SU2C/PCF PCa cohort. Significance determined using student’s t-test.

**Table 1.** Association of six CIN signature gene sets with survival in three PRAD cancer datasets.

|  | **CIN9** | **CIN70** | **CIN25** | **CIN23** | **CIN7** | **CA20** |
| --- | --- | --- | --- | --- | --- | --- |
| **p-value of the Cox**<br>**proportional hazards**<br>**test** | **CN-sig3**<br>**associated**<br>**genes** | **Carter SL**<br>**et al.**<br>**Nat.Genet**<br>**2006** | **Carter SL**<br>**et al.**<br>**Nat.Genet**<br>**2006** | **Bakhoun**<br>**SF et al.**<br>**Nature**<br>**2018** | **Miller**<br>**ET et**<br>**al. BMC**<br>**Cancer**<br>**2020** | **Ogden**<br>**A et**<br>**al.Sci**<br>**Rep**<br>**2017** |
| DKFZ, Cancer Cell<br>2018 BCR-free survival | 0.00012 | 0.00031 | <0.0001 | 0.0083 | 0.00037 | <0.0001 |
| MSK, Cancer Cell 2010<br>Disease-free survival | 0.0068 | 0.88 | 0.0028 | 0.021 | <0.0001 | 0.14 |
| SU2C/PCF DREAM<br>TEAM, PNAS 2019<br>Overall survival | 0.046 | 0.23 | 0.2 | 1 | 0.084 | 0.18 |

## DISCUSSION

PCa evolution is characterized by a few bursts of CIN that fuels genomic heterogeneity and drives cells along an oncogenic path, as opposed to a slow and steady buildup of mutations over time.^5,9,60^ We sought to model the punctuated progression of PCa using immortalized prostate cells to track molecular changes from non-tumorigenic cells to cells isolated from xenograft tumors. In our in vitro model, cells with transient centrosome elimination acquired apparently random CNVs in early progenitors due to chromosomal instability (CIN). Cell populations that then produced xenograft tumors were composed of dominant clones carrying recurrent focal CNVs. Similarly, aggressive subclones that arise in the heterogeneous primary prostate tumor, then migrate to the metastatic niche where these subclones grow to create more genomically and phenotypically homogenous metastatic tumors. Interestingly, our findings suggest that the CIN incurred by centrosome loss did not directly produce a tumorigenic line but, instead, produced a line (CTN9) with an enhanced rate of CIN that eventually gave rise to a line with an oncogenic CNV pattern (CTN9-Line7). What specific genomic alteration(s) caused the increased CIN rate is unknown but, we did observe an increase in centrosome numbers in CTN9, indicating the possible dysregulation of genes involved in centrosome biogenesis. Nevertheless, we identified PCa patient samples with a similar pattern of focal CNVs that defined a unique centrosome loss-associated CNV signature. This CNV signature (CNV-signature 3) is significantly associated with recurrent CNV regions, increased CIN, tumor sample stratification, and poor clinical outcomes. Centrosome loss-associated CIN signature genes also show strong predictive value at both molecular and clinical levels.

Compared to the overproduction of centrosomes (i.e., amplification), centrosome loss has received much less attention in cancer research. However, a growing number of studies have shown centrosome loss in human primary tumor tissues and tumor cell lines.^32,41,78^ Our previous work demonstrated that centrosome loss is prevalent in localized primary prostate carcinoma, and that its frequency is positively associated with tumor grade. Additionally, a high-throughput centrosome staining effort reported that centrosome loss events were more frequent than centrosome amplification in primary human epithelial ovarian cancers (EOCs) from a large cohort sample ^41^. To further understand the impact of centrosome loss on CIN, and tumor progression in general, it will be important to determine how widespread centrosome loss is in different cancers and, when detected, WGS/shallow WGS should be applied. Undoubtedly, the cellular responses to centrosome loss in real human tumor samples will show considerable differences compared to the artificial perturbations in our experimental model.

Regarding types of CIN caused by centrosome loss, it is unclear whether the genomic rearrangements are stochastic events. The structural variations (SV) identified in the centrosome loss samples suggest that SV clusters are located at or near centromere regions, even after applying filters to exclude potential false-positive SVs that are tripped by highly repetitive centromere regions. Indeed, the apparent random and decreased microtubule nucleation capacity around mitotic chromosomes increases the risk of merotelic kinetochore attachments during acentrosomal spindle assembly, making the centromere a susceptible region for breakpoints.^49,50^ Moreover, a pan-cancer CNVs study has shown a fourfold enrichment of breakpoint density in centromeres relative to within chromosome arms; notably, breakpoint number did not correlate with centromere length^48^. It remains an open question as to whether mitotic errors, including the effects of centrosome amplification and loss, affect certain chromosomes and/or distinct chromosomal regions disproportionally. It would be beneficial to have more WGS data on specific CIN etiologies, since all currently available CIN signatures are transcriptional, established retrospectively, and derived from cells with unknown genomic profiles.^79^

To extend the application of the genomic CNV signature, we also identified a set of nine differentially-expressed genes associated with CIN and potentially linked to centrosome loss. Among the CIN9 genes are *KIF11*, a bipolar kinesin-5 motor (aka, Eg5) that can crossbridge and slide antiparallel spindle microtubules apart, thereby powering centrosome separation during mitosis. Eg5 inhibition causes monopolar spindle formation ^70,71^, which resembles an early forming acentrosomal spindle and increases merotelic attachments.^49,50^ Another key gene, *PPP6R3*, is a regulatory subunit of protein phosphatase 6 (PP6) which dephosphorylates Aurora-A, a kinase essential for mitotic progression. Deletion of PPP6R3 results in hyperactivation of Aurora-A ^80^, which plays several important mitotic roles, including centrosome maturation and supernumerary centrosome clustering.^81^ *SMC2* is a core component of the condensin complex that functions in chromosome condensation and the maintenance of chromosomal integrity during mitosis.^82^ Together, these genes highlight the intricate relationships between centrosome biology, cell division, and genomic stability. Future studies should focus on examining expression levels of these genes throughout PCa progression, especially in relation to the timing of centrosome loss, which is still yet to be determined.

One key limitation of this study is the relatively limited sample size used for the analysis of transient centrosome loss samples. The constrained number of available samples may limit the robustness of our conclusions regarding their impact on CIN and tumorigenesis. Future studies with larger cohorts and targeted functional assays will be essential to validate these findings and explore their broader implications. Nevertheless, our multi-omics approach, combined with in vitro cell and tumor models, suggests a punctuated evolutionary path for how the PCa genome is shaped. Centrosome loss-induced CIN signatures have the potential to serve as biomarkers to stratify patient outcomes and guide appropriate treatments.

## EXPERIMENTAL MODEL

### Cell culture

Immortalized human PrEC cells (W. Hahn, MIT) and the derived transient centrosome loss cell lines were cultured at 37°C under 5% CO_2_^32^. Cells were cultured in Iscove’s Modified Dulbecco’s Medium (IMDM) (Corning, cat #15-016-CV) containing 10% fetal bovine serum (FBS), 10 U/ml penicillin, and 10 mg/ml streptomycin (GIBCO, cat: #35050061).

### Animal studies

Mouse experiments were conducted with animal care and committee approval using the Experimental Mouse Shared Resource. 5 x 10^6^ of CTN9-line 2, 4, 5, 7 cells were prepared in 0.1 ml of sterile saline and implanted on both flank of male (NOD/SCID/IL2Rγ^null^) NSG mice (Jackson Laboratories) subcutaneously. Tumors growth was measured twice a week, and tumor volume was estimated according to the formula: [(width)^2^ x length]/2. Animals were sacrificed by CO_2_ when total tumor reached 2000 mm^3^.

## METHODS DETAILS

### PCR

Total DNA was isolated from CTN9 expanded cell lines by using DNeasy Blood & Tissue Kit (Qiagen, cat: #69504) following the manufacture’s guidance. Phusion high-fidelity DNA polymerase (ThermoFisher, cat: #F630S), 5x high-fidelity buffer (ThermoFisher, cat: #F518L), dNTPs (ThermoFisher, cat: #R0181) and DMSO were used for making PCR reactions. Primers were designed to target genes spanning both the short and long arms of the male-specific euchromatic (MSY) region: *RPS4Y1* primers: 5’-ACTTCCTGCGGTTTACGACA-3’ (forward), 5’-GAAGAGACCACGCAAAGCAC-3’ (reverse). *PRKY*: 5’-CTTGGGGACAACATCTGAAGGG-3’ (forward), 5’-TGTGAACTGTGGGAGGTGTATTT-3’ (reverse). *UTY*: 5’-GAGGCAAAGAAAATGGCGGA-3’ (forward), 5’-AATAAATACTGGCTGGGCGGT-3’(reverse). *KDM5D*: 5’-GGCTGTCGTCGTGAACAACT-3’(forward), 5’-TGCCTCCCAATCTTACCTCCA-3’(reverse). *DAZ4*: 5’-TGTCAAAACTGGCGTTCCCT-3’ (foward), 5’-GGCTTCGAGTGGTCAAAGGA-3’ (reverse). Positive control *CEP215*: 5’- TCCTCCTCAGAGAGTTGTTTCTGCC -3’ (forward), 5’- ATCAGAGGGTAGTGGGAAGCCGC -3’ (reverse).

### Fluorescent in situ hybridization (FISH)

2.5 x 10^4^ – 5 x 10^4^ CTN9 expanded cells were plate on a chamber slide (one type of cells per grid) (ThermoFisher, cat: #12-565-8) and cultured at 37°C in a 5% CO_2_ incubator for 24-48 hours until they reached 80-90% confluency. Slide were placed in fixation buffer (75% methanol, 25% acetic acid) for 30 minutes and then warmed to room temperature. Next, slides were treated with denature buffer (70% formamide, 2X SSC in distilled water, added acid to adjust to pH 7.0 and warmed to 73°C for 30 min before use). Slides were placed in 70%, 85%, and 100% ethanol for 2 minutes each and warmed at 37°C until the ethanol evaporated. Denatured Y chromosome FISH probe (Empire Genomics, cat: #CHRY-10-GR) mixture (3 μl of probe was mixed with 12 μl of hybridization buffer per reaction and prewarmed to at 73°C for 5 minutes) was placed on ice for 2 minutes. The probe was briefly centrifuged, placed at 37°C for 15 minutes, and 15 μl of denatured probe added to the sample. Slides were place in a sealed humidity chamber and incubated at 37°C for 16 hours. Next, slides were washed in Buffer1 (0.3% NP-40, 0.4X SSC, pH 7.0), then Buffer 2 (0.1% NP-40, 2X SSC, pH 7.0.), and placed at room temperature for 1 minutes in the dark. Lastly, 15 μl Antifade (ThermoFisher Scientific, cat: #P36935) mounting media with DAPI was applied. Specimens were imaged with a DeltaVision Core system equipped with a microscope (IX71; Olympus) equipped with a 100× objective lens (NA 1.4) and a CoolSNAP HQ2 cooled-CCD camera (Photometrics). Images were acquired with SoftWoRx v1.2 software (Applied Precision).

### Immunofluorescence microscopy

Cells were cultured in 8-well Lab-Tek chamber slides (ThermoFisher, cat: #12-565-8), washed with phosphate-buffered saline (PBS), and fixed in ice-cold methanol for 10 minutes. Following fixation, cells were rehydrated in PBS for 5 minutes and permeabilized with Buffer-A (PBS + 0.5% Triton X-100) for 5 minutes. Non-specific binding was blocked by incubating the cells in Buffer-B (PBS + 5% normal goat serum and 0.1% Triton X-100) for 30 minutes at room temperature. Cells were then incubated with primary antibodies diluted in Buffer-B for 1 hour at room temperature. After three washes with Buffer-A, cells were incubated with secondary antibodies and Hoechst 33342 for 1 hour at room temperature. Following three additional washes with Buffer-A (5 minutes each), cells were mounted using a medium composed of 0.1 M n-propyl gallate, PBS, and 90% glycerol (by volume). For immunostaining, the following primary antibodies were used: anti-CEP135 (Abcam, cat: #ab75005; 1:500 for cells, 1:100 for tissue) to label centrioles and anti-γtubulin (Sigma Aldrich, cat: #GTU-88; 1:500) to stain pericentriolar material. Specimens were imaged as described in the FISH method.

### Molecular biology and WGS processing pipeline

Total DNA was isolated from cultured cells using DNeasy Blood & Tissue Kit (Qiagen, cat: #69504). The isolated DNA was subject to DNA electrophoresis on 1% agarose gels to exempt DNA degradation and RNA contamination. Total DNA was measured using Qubit DNA Assay kit in Qubit 2.0 fluorometer (Life Technologies, cat: #Q33230). The genomic DNA was randomly fragmented by sonication to the size of 350 bp, DNA fragments were end polished, A-tailed, and ligated with the full-length adapters of Illumina sequencing followed by further PCR amplification and qPCR QC analyses. The product was sequenced on Illumina NovaSeq 6000 platform with average sequencing depth at least 30x of all samples; average coverage of genome was at least 4x at 99.6%, 10x at 98.0% and 20x at 83.7%. WGS sequencing raw data was checked for quality using FastQC and DNA-seq adapters were trimmed using Trimmomatic-0.39 ^84^. Then, Burrows-Wheeler Aligner (BWA/0.7.17) MEM ^85^ was utilized to map the paired-end trimmed reads to the human reference genome “GRCh38.d1.vd1_BWA.tar.gz” downloaded from https://gdc.cancer.gov/about-data/gdc-data-processing/gdc-reference-files. Samtools/1.19.2 ^86^ was then used to convert the aligned files from .sam to .bam files and then sorted.

### Somatic structural variation for WGS

Somatic structural variations (SV) were called from pre-processed WGS data by utilizing DELLY/0.9.1 ^42^ with tumor-normal paired mode used with PrEC-Hahn parental as normal and transient centrosome loss cells and tumor tissue as tumors.

### Somatic copy number variation for WGS

Somatic copy number variation (CNV) was called from pre-processed WGS data by utilizing CNVkit/0.9.10 ^43^ for non-allelic specific CNVs, and Sequenza ^54^ and FACETS ^63^ for allelic specific CNVs. Tumor-normal paired pipelines were used with PrEC-Hahn parental as normal and transient centrosome loss cells and tumor tissue as tumors.

### Tumor purity score

The estimated tumor purity of centrosome loss samples was calculated through FACETS from tumor-normal pairs with initial seed set. Purity represents the fraction of cancer cells in the tumor sample, whereas in the transient centrosome loss sample, it represents the cellular prevalence of focal CNVs.

### CNV signature extraction

We used preprocessed CNV data from either WES or WGS data of PCa patient samples from multiple datasets^2,3,23,55–57^. Absolute CNVs were called from tumor-normal paired samples by utilizing Sequenza or FACETS. To identify CNV signatures, we used the Sigminer^55^ algorithm where 80 copy number related genomic alteration components were extracted from the CNV data. Thus, a sample-by-components matrix can be created, which was treated as the input to the non-negative matrix factorization (NMF) algorithm for extracting signatures. The output matrices from NMF provides the relative exposure of each signature for each tumor, as well as the relative weight of each copy number feature in each signature. Samples were classified by the signature for which they had the maximum exposure value.

### RNA isolation and sequencing

Total RNA was isolated from cells in triplicate using the RNeasy mini kit (Qiagen, cat: #74104). Isolated RNA was subjected to quality control using an Agilent 2100 bioanalyzer. RNA samples that passed quality control were followed by library preparation and sequencing on the Illumina NovaSeq 6000 platform.

### RNA-seq processing pipeline

Raw reads were first passed through FastQC, adapters were trimmed using Trimmomatic-0.39, reads were aligned with the reference genome “GDC.h38.d1.vd1 STAR2 Index Files (v36)” (downloaded from https://gdc.cancer.gov/about-data/gdc-data-processing/gdc-reference-files using STAR/2.7.10a), and counts were quantified using the Rsubread R package. Raw data and counts matrix can be found in the Gene Expression Omnibus (GEO) database (accession number: GSE250174). The edgeR pipeline using TMM normalization and voom with default parameters was used to normalize the counts matrix. To filter out low-expressing genes, we performed calculations of counts per million (cpm) using the raw read counts matrix. Genes with less than half of the samples exceeding 1 cpm were subsequently removed.

### Differential expression analysis

Differentially expressed genes were identified for RNA-seq data using the limma R package. P-value were adjusted for multiple testing using the Benjamini-Hochberg method.

### scRNA-seq data analyses

Pair-matched radical prostatectomy scRNA-seq data^52^ were downloaded from the GEO database (accession number: GSE176031). The R package Seurat/5.1.0 was used for the subsequent data processing. Read count matrices were filtered to remove low quality cells with a filter defined as greater than 5% mitochondria transcripts and fewer than 200 genes detected. The R package SingleR/1.0 ^87^ was used to identify broad lineages such as epithelial cells, stroma cells, and immune cells. Copy number analysis was performed on the epithelial cell populations from all four patients with matched tumor and normal sample pairs, utilizing R package InferCNV/1.19.1 with cutoff = 0.1 (inferCNV from the Trinity CTAT Project: https://github.com/broadinstitute/inferCNV). Uphyloplot2/2.3 ^53^ was then used to depict the clonality and evolution information based on the inferCNV results in epithelial cell populations from individual patients’ tumor and normal samples with threshold set as at least 5% of cells to be included for plotting.

### Signature gene selection

TCGA PRAD mRNA raw read count data for all patients were acquired from the GDC data portal. To filter out low-expressing genes, we calculated counts per million (cpm) using the raw read counts matrix and removed genes with less than half of the samples exceeding 1 cpm. The edgeR pipeline using TMM normalization and voom with default parameters was used to normalize the counts matrix. Patient samples were then labeled based on CNV-signature classification: TCGA-PRAD patients classified as CNV-signature 3 were assigned a binary label of “1” (CNV-signature 3 -positive), while all remaining samples were labeled “0” (CNV-signature 3 -negative). Using these binary labels as the outcome variable, we applied least absolute shrinkage and selection operator (Lasso) logistic regression to identify signature genes predictive of CNV-signature 3 status. To optimize the regularization parameter (λ), we implemented 10-fold cross-validation: the dataset was randomly partitioned into 10 equally sized folds, and the model was iteratively trained on 9 folds and validated on the withheld fold. The optimal λ (λ_min) was selected as the value minimizing the cross-validated binomial deviance. After selecting λ_min, the final Lasso model was retrained on the entire dataset using this parameter. Genes with non-zero coefficients in the retrained model were extracted as the CNV-signature 3 -associated signature genes. The above analyses were performed using the R package glmnet (version 4.1-8). The robustness of the identified signature genes was validated in three different independent PCa datasets. ^47,72,83^

### Statistics

R/4.2.2 and GraphPad Prism/9.0.2 were used in all statistical tests for computational and in vitro analyses. Specific statistical methods and details are described in main and supplemental figure legends.

### Study approval

This study was approved by the IACUC for the care and use of laboratory animals at the University of Arizona (protocol #2021 - 0772)

## Data availability

The data generated in this study are publicly available in Gene Expression Omnibus (GEO) under accession number GSE250174 and Sequence Read Archive (SRA) under the project ID: PRJNA1053425. Previously published data analyzed in this study were obtained from GEO under accession number GSE176031 and cBioportal: Metastatic prostate adenocarcinoma (SU2C/PCFD Dream Team, PNAS 2019: https://cbioportal-datahub.s3.amazonaws.com/prad_su2c_2019.tar.gz), Prostate Cancer (DKFZ, Cancer Cell 2018: https://cbioportal-datahub.s3.amazonaws.com/prostate_dkfz_2018.tar.gz), and Prostate Adenocarcinoma (MSK, Cancer Cell 2010: https://cbioportal-datahub.s3.amazonaws.com/prad_mskcc.tar.gz). FACETS and Sequenza somatic CNV calls from PCa cohorts were obtained from Wang et al (https://github.com/XSLiuLab/PC_CNA_signature). ^55^ All code and processed data are available on Github (https://github.com/JiawenYang16/centrosome_loss_and_PCa).

## ACKNOWLEDGMENTS

This work was supported by National Cancer Institute (NCI) of the National Institutes of Health (NIH) grant (R01CA242226 to G.C.R. and A.E.C., and R01CA251729 to M.P., and T32CA009213 to M.R.C.). We thank the Experimental Mouse Shared Resource (ESMR) at the University of Arizona Cancer Center (UACC) supported by the NCI of the NIH under award P30CA023074. We thank the members of Rogers and Padi labs for their help with data interpretation, development, and discussions. We thank Drs. Guang Yao, Ryan Gutenkunst, and Andrew Paek at the University of Arizona for insightful suggestions.

## AUTHOR CONTRIBUTIONS

Conceptualization: G.C.R., M.P., J.Y. and J.M.R.; Methodology: J.Y., D.D.O.P., M.R.C. and C.C.; Experiments, J.Y, M.R.C., E.L., and M.W.; Data analysis, J.Y., M.W., D.W.B., and C.C.; Resources, G.C.R., A.E.C., and M.P.; Writing – initial draft, J.Y.; Review and editing, G.C.R., M.P., J.Y., and J.M.R.; Supervision, G.C.R., M.P.; Funding acquisition, G.C.R., A.E.C., and M.P.

## DECLARATION OF INTERESTS

The authors declare no competing interests.

## Notes

### Competing Interest Statement

The authors have declared no competing interest.

## REFERENCES

1. Rebello, R.J., Oing, C., Knudsen, K.E., Loeb, S., Johnson, D.C., Reiter, R.E., Gillessen, S., Kwast, T.V. der, and Bristow, R.G. (2021). Prostate cancer. Nat. Rev. Dis. Prim. 7, 9. 10.1038/s41572-020-00243-0.

2. Network, T.C.G.A.R., Abeshouse, A., Ahn, J., Akbani, R., Ally, A., Amin, S., Andry, C.D., Annala, M., Aprikian, A., Armenia, J., et al. (2015). The Molecular Taxonomy of Primary Prostate Cancer. Cell 163, 1011–1025. 10.1016/j.cell.2015.10.025.

3. Berger, M.F., Lawrence, M.S., Demichelis, F., Drier, Y., Cibulskis, K., Sivachenko, A.Y., Sboner, A., Esgueva, R., Pflueger, D., Sougnez, C., et al. (2011). The genomic complexity of primary human prostate cancer. Nature 470, 214–220. 10.1038/nature09744.

4. Hieronymus, H., Schultz, N., Gopalan, A., Carver, B.S., Chang, M.T., Xiao, Y., Heguy, A., Huberman, K., Bernstein, M., Assel, M., et al. (2014). Copy number alteration burden predicts prostate cancer relapse. Proc National Acad Sci 111, 11139–11144. 10.1073/pnas.1411446111.

5. Baca, S.C., Prandi, D., Lawrence, M.S., Mosquera, J.M., Romanel, A., Drier, Y., Park, K., Kitabayashi, N., MacDonald, T.Y., Ghandi, M., et al. (2013). Punctuated Evolution of Prostate Cancer Genomes. Cell 153, 666–677. 10.1016/j.cell.2013.03.021.

6. Peng, Y., Song, Y., and Wang, H. (2022). Systematic Elucidation of the Aneuploidy Landscape and Identification of Aneuploidy Driver Genes in Prostate Cancer. Front. Cell Dev. Biol. 9, 723466. 10.3389/fcell.2021.723466.

7. Stopsack, K.H., Whittaker, C.A., Gerke, T.A., Loda, M., Kantoff, P.W., Mucci, L.A., and Amon, A. (2019). Aneuploidy drives lethal progression in prostate cancer. Proc. Natl. Acad. Sci. 116, 11390–11395. 10.1073/pnas.1902645116.

8. Guan, Y., Wang, X., Guan, K., Wang, D., Bi, X., Xiao, Z., Xiao, Z., Shan, X., Hu, L., Ma, J., et al. (2022). Copy number variation of urine exfoliated cells by low-coverage whole genome sequencing for diagnosis of prostate adenocarcinoma: a prospective cohort study. BMC Méd. Genom. 15, 104. 10.1186/s12920-022-01253-5.

9. Shen, M.M. (2013). Chromoplexy: A New Category of Complex Rearrangements in the Cancer Genome. Cancer Cell 23, 567–569. 10.1016/j.ccr.2013.04.025.

10. Tomlins, S.A., Laxman, B., Dhanasekaran, S.M., Helgeson, B.E., Cao, X., Morris, D.S., Menon, A., Jing, X., Cao, Q., Han, B., et al. (2007). Distinct classes of chromosomal rearrangements create oncogenic ETS gene fusions in prostate cancer. Nature 448, 595–599. 10.1038/nature06024.

11. Shen, M.M., and Abate-Shen, C. (2010). Molecular genetics of prostate cancer: new prospects for old challenges. Gene Dev 24, 1967–2000. 10.1101/gad.1965810.

12. Bakhoum, S.F., and Cantley, L.C. (2018). The Multifaceted Role of Chromosomal Instability in Cancer and Its Microenvironment. Cell 174, 1347–1360. 10.1016/j.cell.2018.08.027.

13. Armenia, J., Wankowicz, S.A.M., Liu, D., Gao, J., Kundra, R., Reznik, E., Chatila, W.K., Chakravarty, D., Han, G.C., Coleman, I., et al. (2018). The long tail of oncogenic drivers in prostate cancer. Nat Genet 50, 645–651. 10.1038/s41588-018-0078-z.

14. Boutros, P.C., Fraser, M., Harding, N.J., Borja, R. de, Trudel, D., Lalonde, E., Meng, A., Hennings-Yeomans, P.H., McPherson, A., Sabelnykova, V.Y., et al. (2015). Spatial genomic heterogeneity within localized, multifocal prostate cancer. Nat Genet 47, 736–745. 10.1038/ng.3315.

15. Ben-David, U., and Amon, A. (2020). Context is everything: aneuploidy in cancer. Nat Rev Genet 21, 44–62. 10.1038/s41576-019-0171-x.

16. Bostwick, D.G., and Qian, J. (2004). High-grade prostatic intraepithelial neoplasia. Mod. Pathol. 17, 360–379. 10.1038/modpathol.3800053.

17. Bostwick, D.G., Shan, A., Qian, J., Darson, M., Maihle, N.J., Jenkins, R.B., and Cheng, L. (1998). Independent origin of multiple foci of prostatic intraepithelial neoplasia: comparison with matched foci of prostate carcinoma. Cancer 83, 1995–2002. 10.1002/(sici)1097-0142(19981101)83:9<1995::aid-cncr16>3.0.co;2-2.

18. Qian, J., Jenkins, R.B., and Bostwick, D.G. (1997). Detection of chromosomal anomalies and c-myc gene amplification in the cribriform pattern of prostatic intraepithelial neoplasia and carcinoma by fluorescence in situ hybridization. Mod. Pathol. : Off. J. United States Can. Acad. Pathol., Inc 10, 1113–1119.

19. Myers, R.B., Brown, D., Oelschlager, D.K., Waterbor, J.W., Marshall, M.E., Srivastava, S., Stockard, C.R., Urban, D.A., and Grizzle, W.E. (1996). Elevated serum levels of p105erbB-2 in patients with advanced-stage prostatic adenocarcinoma. Int. J. Cancer 69, 398–402. 10.1002/(sici)1097-0215(19961021)69:5<398::aid-ijc8>3.0.co;2-0.

20. Emmert-Buck, M.R., Vocke, C.D., Pozzatti, R.O., Duray, P.H., Jennings, S.B., Florence, C.D., Zhuang, Z., Bostwick, D.G., Liotta, L.A., and Linehan, W.M. (1995). Allelic loss on chromosome 8p12-21 in microdissected prostatic intraepithelial neoplasia. Cancer Res. 55, 2959–2962.

21. Tomlins, S.A., Rhodes, D.R., Perner, S., Dhanasekaran, S.M., Mehra, R., Sun, X.-W., Varambally, S., Cao, X., Tchinda, J., Kuefer, R., et al. (2005). Recurrent Fusion of *TMPRSS2* and ETS Transcription Factor Genes in Prostate Cancer. Science 310, 644–648. 10.1126/science.1117679.

22. Nguyen, B., Fong, C., Luthra, A., Smith, S.A., DiNatale, R.G., Nandakumar, S., Walch, H., Chatila, W.K., Madupuri, R., Kundra, R., et al. (2022). Genomic characterization of metastatic patterns from prospective clinical sequencing of 25,000 patients. Cell 185, 563–575.e11. 10.1016/j.cell.2022.01.003.

23. Robinson, D., Van Allen, E.M., Wu, Y.-M., Schultz, N., Lonigro, R.J., Mosquera, J.-M., Montgomery, B., Taplin, M.-E., Pritchard, C.C., Attard, G., et al. (2015). Integrative Clinical Genomics of Advanced Prostate Cancer. Cell 161, 1215–1228. 10.1016/j.cell.2015.05.001.

24. Mateo, J., Seed, G., Bertan, C., Rescigno, P., Dolling, D., Figueiredo, I., Miranda, S., Rodrigues, D.N., Gurel, B., Clarke, M., et al. (2019). Genomics of lethal prostate cancer at diagnosis and castration-resistance. J. Clin. Investig. 130, 1743–1751. 10.1172/jci132031.

25. Stopsack, K.H., Nandakumar, S., Wibmer, A.G., Haywood, S., Weg, E.S., Barnett, E.S., Kim, C.J., Carbone, E.A., Vasselman, S.E., Nguyen, B., et al. (2020). Oncogenic Genomic Alterations, Clinical Phenotypes, and Outcomes in Metastatic Castration-Sensitive Prostate Cancer. Clin. Cancer Res. 26, 3230–3238. 10.1158/1078-0432.ccr-20-0168.

26. Gao, R., Davis, A., McDonald, T.O., Sei, E., Shi, X., Wang, Y., Tsai, P.-C., Casasent, A., Waters, J., Zhang, H., et al. (2016). Punctuated copy number evolution and clonal stasis in triple-negative breast cancer. Nat. Genet. 48, 1119–1130. 10.1038/ng.3641.

27. Cooper, C.S., Eeles, R., Wedge, D.C., Loo, P.V., Gundem, G., Alexandrov, L.B., Kremeyer, B., Butler, A., Lynch, A.G., Camacho, N., et al. (2015). Analysis of the genetic phylogeny of multifocal prostate cancer identifies multiple independent clonal expansions in neoplastic and morphologically normal prostate tissue. Nat. Genet. 47, 367–372. 10.1038/ng.3221.

28. Hong, M.K.H., Macintyre, G., Wedge, D.C., Loo, P.V., Patel, K., Lunke, S., Alexandrov, L.B., Sloggett, C., Cmero, M., Marass, F., et al. (2015). Tracking the origins and drivers of subclonal metastatic expansion in prostate cancer. Nat. Commun. 6, 6605. 10.1038/ncomms7605.

29. Levine, M.S., and Holland, A.J. (2018). The impact of mitotic errors on cell proliferation and tumorigenesis. Genes Dev. 32, 620–638. 10.1101/gad.314351.118.

30. Nigg, E.A. (2006). Origins and consequences of centrosome aberrations in human cancers. Int. J. Cancer 119, 2717–2723. 10.1002/ijc.22245.

31. Nigg, E.A., and Holland, A.J. (2018). Once and only once: mechanisms of centriole duplication and their deregulation in disease. Nat. Rev. Mol. Cell Biol. 19, 297–312. 10.1038/nrm.2017.127.

32. Wang, M., Nagle, R.B., Knudsen, B.S., Cress, A.E., and Rogers, G.C. (2020). Centrosome loss results in an unstable genome and malignant prostate tumors. Oncogene 39, 399–413. 10.1038/s41388-019-0995-z.

33. Khodjakov, A., and Rieder, C.L. (2001). Centrosomes Enhance the Fidelity of Cytokinesis in Vertebrates and Are Required for Cell Cycle Progression. J. Cell Biol. 153, 237–242. 10.1083/jcb.153.1.237.

34. Sir, J.-H., Pütz, M., Daly, O., Morrison, C.G., Dunning, M., Kilmartin, J.V., and Gergely, F. (2013). Loss of centrioles causes chromosomal instability in vertebrate somatic cellsCentrosomes maintain chromosome stability. J Cell Biology 203, 747–756. 10.1083/jcb.201309038.

35. Lambrus, B.G., Uetake, Y., Clutario, K.M., Daggubati, V., Snyder, M., Sluder, G., and Holland, A.J. (2015). p53 protects against genome instability following centriole duplication failure. J. Cell Biol. 210, 63–77. 10.1083/jcb.201502089.

36. Wong, Y.L., Anzola, J.V., Davis, R.L., Yoon, M., Motamedi, A., Kroll, A., Seo, C.P., Hsia, J.E., Kim, S.K., Mitchell, J.W., et al. (2015). Reversible centriole depletion with an inhibitor of Polo-like kinase 4. Science 348, 1155–1160. 10.1126/science.aaa5111.

37. Silkworth, W.T., Nardi, I.K., Scholl, L.M., and Cimini, D. (2009). Multipolar Spindle Pole Coalescence Is a Major Source of Kinetochore Mis-Attachment and Chromosome Mis-Segregation in Cancer Cells. PLoS ONE 4, e6564. 10.1371/journal.pone.0006564.

38. Ganem, N.J., Godinho, S.A., and Pellman, D. (2009). A mechanism linking extra centrosomes to chromosomal instability. Nature 460, 278–282. 10.1038/nature08136.

39. Chan, J.Y. (2011). A Clinical Overview of Centrosome Amplification in Human Cancers. Int. J. Biol. Sci. 7, 1122–1144. 10.7150/ijbs.7.1122.

40. Cosenza, M.R., and Krämer, A. (2016). Centrosome amplification, chromosomal instability and cancer: mechanistic, clinical and therapeutic issues. Chromosom. Res. 24, 105–126. 10.1007/s10577-015-9505-5.

41. Morretton, J., Simon, A., Herbette, A., Barbazan, J., Pérez-González, C., Cosson, C., Mboup, B., Latouche, A., Popova, T., Kieffer, Y., et al. (2022). A catalog of numerical centrosome defects in epithelial ovarian cancers. EMBO Mol. Med. 14, e15670. 10.15252/emmm.202215670.

42. Rausch, T., Zichner, T., Schlattl, A., Stütz, A.M., Benes, V., and Korbel, J.O. (2012). DELLY: structural variant discovery by integrated paired-end and split-read analysis. Bioinformatics 28, i333–i339. 10.1093/bioinformatics/bts378.

43. Talevich, E., Shain, A.H., Botton, T., and Bastian, B.C. (2016). CNVkit: Genome-Wide Copy Number Detection and Visualization from Targeted DNA Sequencing. PLoS Comput. Biol. 12, e1004873. 10.1371/journal.pcbi.1004873.

44. Macoska, J.A., Trybus, T.M., Benson, P.D., Sakr, W.A., Grignon, D.J., Wojno, K.D., Pietruk, T., and Powell, I.J. (1995). Evidence for three tumor suppressor gene loci on chromosome 8p in human prostate cancer. Cancer Res. 55, 5390–5395.

45. Bostwick, D.G., and Cheng, L. (2020). Urologic Surgical Pathology. 415–525.e42. 10.1016/b978-0-323-54941-7.00009-8.

46. Dessel, L.F. van, Riet, J. van, Smits, M., Zhu, Y., Hamberg, P., Heijden, M.S. van der, Bergman, A.M., Oort, I.M. van, Wit, R. de, Voest, E.E., et al. (2019). The genomic landscape of metastatic castration-resistant prostate cancers reveals multiple distinct genotypes with potential clinical impact. Nat Commun 10, 5251. 10.1038/s41467-019-13084-7.

47. Gerhauser, C., Favero, F., Risch, T., Simon, R., Feuerbach, L., Assenov, Y., Heckmann, D., Sidiropoulos, N., Waszak, S.M., Hübschmann, D., et al. (2018). Molecular Evolution of Early-Onset Prostate Cancer Identifies Molecular Risk Markers and Clinical Trajectories. Cancer Cell 34, 996–1011.e8. 10.1016/j.ccell.2018.10.016.

48. Shih, J., Sarmashghi, S., Zhakula-Kostadinova, N., Zhang, S., Georgis, Y., Hoyt, S.H., Cuoco, M.S., Gao, G.F., Spurr, L.F., Berger, A.C., et al. (2023). Cancer aneuploidies are shaped primarily by effects on tumour fitness. Nature 619, 793–800. 10.1038/s41586-023-06266-3.

49. Cimini, D., Howell, B., Maddox, P., Khodjakov, A., Degrassi, F., and Salmon, E.D. (2001). Merotelic Kinetochore Orientation Is a Major Mechanism of Aneuploidy in Mitotic Mammalian Tissue Cells. J. Cell Biol. 153, 517–528. 10.1083/jcb.153.3.517.

50. Gregan, J., Polakova, S., Zhang, L., Tolić-Nørrelykke, I.M., and Cimini, D. (2011). Merotelic kinetochore attachment: causes and effects. Trends Cell Biol 21, 374–381. 10.1016/j.tcb.2011.01.003.

51. Chen, X., Agustinus, A.S., Li, J., DiBona, M., and Bakhoum, S.F. (2025). Chromosomal instability as a driver of cancer progression. Nat. Rev. Genet. 26, 31–46. 10.1038/s41576-024-00761-7.

52. Song, H., Weinstein, H.N.W., Allegakoen, P., Wadsworth, M.H., Xie, J., Yang, H., Castro, E.A., Lu, K.L., Stohr, B.A., Feng, F.Y., et al. (2022). Single-cell analysis of human primary prostate cancer reveals the heterogeneity of tumor-associated epithelial cell states. Nat Commun 13, 141. 10.1038/s41467-021-27322-4.

53. Kurtenbach, S., Cruz, A.M., Rodriguez, D.A., Durante, M.A., and Harbour, J.W. (2021). Uphyloplot2: visualizing phylogenetic trees from single-cell RNA-seq data. BMC Genom. 22, 419. 10.1186/s12864-021-07739-3.

54. Favero, F., Joshi, T., Marquard, A.M., Birkbak, N.J., Krzystanek, M., Li, Q., Szallasi, Z., and Eklund, A.C. (2015). Sequenza: allele-specific copy number and mutation profiles from tumor sequencing data. Ann. Oncol. 26, 64–70. 10.1093/annonc/mdu479.

55. Wang, S., Li, H., Song, M., Tao, Z., Wu, T., He, Z., Zhao, X., Wu, K., and Liu, X.-S. (2021). Copy number signature analysis tool and its application in prostate cancer reveals distinct mutational processes and clinical outcomes. PLoS Genet. 17, e1009557. 10.1371/journal.pgen.1009557.

56. Grasso, C.S., Wu, Y.-M., Robinson, D.R., Cao, X., Dhanasekaran, S.M., Khan, A.P., Quist, M.J., Jing, X., Lonigro, R.J., Brenner, J.C., et al. (2012). The mutational landscape of lethal castration-resistant prostate cancer. Nature 487, 239–243. 10.1038/nature11125.

57. Beltran, H., Rickman, D.S., Park, K., Chae, S.S., Sboner, A., MacDonald, T.Y., Wang, Y., Sheikh, K.L., Terry, S., Tagawa, S.T., et al. (2011). Molecular Characterization of Neuroendocrine Prostate Cancer and Identification of New Drug Targets. Cancer Discov. 1, 487–495. 10.1158/2159-8290.cd-11-0130.

58. Kohli, M., Wang, L., Xie, F., Sicotte, H., Yin, P., Dehm, S.M., Hart, S.N., Vedell, P.T., Barman, P., Qin, R., et al. (2015). Mutational Landscapes of Sequential Prostate Metastases and Matched Patient Derived Xenografts during Enzalutamide Therapy. PLoS ONE 10, e0145176. 10.1371/journal.pone.0145176.

59. Sansregret, L., Vanhaesebroeck, B., and Swanton, C. (2018). Determinants and clinical implications of chromosomal instability in cancer. Nat. Rev. Clin. Oncol. 15, 139–150. 10.1038/nrclinonc.2017.198.

60. Turajlic, S., Sottoriva, A., Graham, T., and Swanton, C. (2019). Resolving genetic heterogeneity in cancer. Nat. Rev. Genet. 20, 404–416. 10.1038/s41576-019-0114-6.

61. Hosea, R., Hillary, S., Naqvi, S., Wu, S., and Kasim, V. (2024). The two sides of chromosomal instability: drivers and brakes in cancer. Signal Transduct. Target. Ther. 9, 75. 10.1038/s41392-024-01767-7.

62. Lukow, D.A., Sausville, E.L., Suri, P., Chunduri, N.K., Wieland, A., Leu, J., Smith, J.C., Girish, V., Kumar, A.A., Kendall, J., et al. (2021). Chromosomal instability accelerates the evolution of resistance to anti-cancer therapies. Dev. Cell 56, 2427–2439.e4. 10.1016/j.devcel.2021.07.009.

63. Shen, R., and Seshan, V.E. (2016). FACETS: allele-specific copy number and clonal heterogeneity analysis tool for high-throughput DNA sequencing. Nucleic Acids Res. 44, e131–e131. 10.1093/nar/gkw520.

64. Mermel, C.H., Schumacher, S.E., Hill, B., Meyerson, M.L., Beroukhim, R., and Getz, G. (2011). GISTIC2.0 facilitates sensitive and confident localization of the targets of focal somatic copy-number alteration in human cancers. Genome Biol 12, R41. 10.1186/gb-2011-12-4-r41.

65. Kautto, E.A., Bonneville, R., Miya, J., Yu, L., Krook, M.A., Reeser, J.W., and Roychowdhury, S. (2016). Performance evaluation for rapid detection of pan-cancer microsatellite instability with MANTIS. Oncotarget 8, 7452–7463. 10.18632/oncotarget.13918.

66. Rodemoyer, B., Kariyawasam, G., Subramanian, V., and Schmidt, K. (2025). Condensin II interacts with BLM helicase in S phase to maintain genome stability. Commun. Biol. 8, 492. 10.1038/s42003-025-07916-0.

67. Dávalos, V., Súarez-López, L., Castaño, J., Messent, A., Abasolo, I., Fernandez, Y., Guerra-Moreno, A., Espín, E., Armengol, M., Musulen, E., et al. (2012). Human SMC2 Protein, a Core Subunit of Human Condensin Complex, Is a Novel Transcriptional Target of the WNT Signaling Pathway and a New Therapeutic Target*. J. Biol. Chem. 287, 43472–43481. 10.1074/jbc.m112.428466.

68. Murakami-Tonami, Y., Kishida, S., Takeuchi, I., Katou, Y., Maris, J.M., Ichikawa, H., Kondo, Y., Sekido, Y., Shirahige, K., Murakami, H., et al. (2014). Inactivation of SMC2 shows a synergistic lethal response in MYCN-amplified neuroblastoma cells. Cell Cycle 13, 1115–1131. 10.4161/cc.27983.

69. Bellelli, R., Borel, V., Logan, C., Svendsen, J., Cox, D.E., Nye, E., Metcalfe, K., O’Connell, S.M., Stamp, G., Flynn, H.R., et al. (2018). Polε Instability Drives Replication Stress, Abnormal Development, and Tumorigenesis. Mol. Cell 70, 707–721.e7. 10.1016/j.molcel.2018.04.008.

70. Slangy, A., Lane, H.A., d’Hérin, P., Harper, M., Kress, M., and Niggt, E.A. (1995). Phosphorylation by p34cdc2 regulates spindle association of human Eg5, a kinesin-related motor essential for bipolar spindle formation in vivo. Cell 83, 1159–1169. 10.1016/0092-8674(95)90142-6.

71. Mayer, T.U., Kapoor, T.M., Haggarty, S.J., King, R.W., Schreiber, S.L., and Mitchison, T.J. (1999). Small Molecule Inhibitor of Mitotic Spindle Bipolarity Identified in a Phenotype-Based Screen. Science 286, 971–974. 10.1126/science.286.5441.971.

72. Taylor, B.S., Schultz, N., Hieronymus, H., Gopalan, A., Xiao, Y., Carver, B.S., Arora, V.K., Kaushik, P., Cerami, E., Reva, B., et al. (2010). Integrative Genomic Profiling of Human Prostate Cancer. Cancer Cell 18, 11–22. 10.1016/j.ccr.2010.05.026.

73. Ross-Adams, H., Lamb, A.D., Dunning, M.J., Halim, S., Lindberg, J., Massie, C.M., Egevad, L.A., Russell, R., Ramos-Montoya, A., Vowler, S.L., et al. (2015). Integration of copy number and transcriptomics provides risk stratification in prostate cancer: A discovery and validation cohort study. EBioMedicine 2, 1133–1144. 10.1016/j.ebiom.2015.07.017.

74. Bakhoum, S.F., Ngo, B., Laughney, A.M., Cavallo, J.-A., Murphy, C.J., Ly, P., Shah, P., Sriram, R.K., Watkins, T.B.K., Taunk, N.K., et al. (2018). Chromosomal instability drives metastasis through a cytosolic DNA response. Nature 553, 467–472. 10.1038/nature25432.

75. Carter, S.L., Eklund, A.C., Kohane, I.S., Harris, L.N., and Szallasi, Z. (2006). A signature of chromosomal instability inferred from gene expression profiles predicts clinical outcome in multiple human cancers. Nat. Genet. 38, 1043–1048. 10.1038/ng1861.

76. Ogden, A., Rida, P.C.G., and Aneja, R. (2017). Prognostic value of CA20, a score based on centrosome amplification-associated genes, in breast tumors. Sci. Rep. 7, 262. 10.1038/s41598-017-00363-w.

77. Miller, E.T., You, S., Cadaneanu, R.M., Kim, M., Yoon, J., Liu, S.T., Li, X., Kwan, L., Hodge, J., Quist, M.J., et al. (2020). Chromosomal instability in untreated primary prostate cancer as an indicator of metastatic potential. BMC Cancer 20, 398. 10.1186/s12885-020-06817-1.

78. Sauer, C.M., Hall, J.A., Couturier, D.-L., Bradley, T., Piskorz, A.M., Griffiths, J., Sawle, A., Eldridge, M.D., Smith, P., Hosking, K., et al. (2023). Molecular landscape and functional characterization of centrosome amplification in ovarian cancer. Nat. Commun. 14, 6505. 10.1038/s41467-023-41840-3.

79. Drews, R.M., Hernando, B., Tarabichi, M., Haase, K., Lesluyes, T., Smith, P.S., Gavarró, L.M., Couturier, D.-L., Liu, L., Schneider, M., et al. (2022). A pan-cancer compendium of chromosomal instability. Nature 606, 976–983. 10.1038/s41586-022-04789-9.

80. Heo, J., Larner, J.M., and Brautigan, D.L. (2020). Protein kinase CK2 phosphorylation of SAPS3 subunit increases PP6 phosphatase activity with Aurora A kinase. Biochem. J. 477, 431–444. 10.1042/bcj20190740.

81. Navarro-Serer, B., Childers, E.P., Hermance, N.M., Mercadante, D., and Manning, A.L. (2019). Aurora A inhibition limits centrosome clustering and promotes mitotic catastrophe in cells with supernumerary centrosomes. Oncotarget 10, 1649–1659. 10.18632/oncotarget.26714.

82. Yokomori, K. (2003). Protein Complexes that Modify Chromatin. Curr. Top. Microbiol. Immunol. 274, 79–112. 10.1007/978-3-642-55747-7_4.

83. Abida, W., Cyrta, J., Heller, G., Prandi, D., Armenia, J., Coleman, I., Cieslik, M., Benelli, M., Robinson, D., Allen, E.M.V., et al. (2019). Genomic correlates of clinical outcome in advanced prostate cancer. Proc. Natl. Acad. Sci. United States Am. 116, 11428–11436. 10.1073/pnas.1902651116.

84. Bolger, A.M., Lohse, M., and Usadel, B. (2014). Trimmomatic: a flexible trimmer for Illumina sequence data. Bioinformatics 30, 2114–2120. 10.1093/bioinformatics/btu170.

85. Li, H., and Durbin, R. (2009). Fast and accurate short read alignment with Burrows–Wheeler transform. Bioinformatics 25, 1754–1760. 10.1093/bioinformatics/btp324.

86. Li, H., Handsaker, B., Wysoker, A., Fennell, T., Ruan, J., Homer, N., Marth, G., Abecasis, G., Durbin, R., and Subgroup, 1000 Genome Project Data Processing (2009). The Sequence Alignment/Map format and SAMtools. Bioinform. (Oxf., Engl.) 25, 2078–2079. 10.1093/bioinformatics/btp352.

87. Aran, D., Looney, A.P., Liu, L., Wu, E., Fong, V., Hsu, A., Chak, S., Naikawadi, R.P., Wolters, P.J., Abate, A.R., et al. (2019). Reference-based analysis of lung single-cell sequencing reveals a transitional profibrotic macrophage. Nat. Immunol. 20, 163–172. 10.1038/s41590-018-0276-y.

