## Supplementary material for "Transient centrosome loss in cultured prostate epithelial cells induces chromosomal instability to produce an oncogenic genotype that correlates with poor clinical outcomes": Figure S, Table S

**Supplemental Figures**

Supplemental Figures 1-12.	p2-22

**Supplemental Tables**

Supplemental Tables 1-5.	p23 - 29

**References**	p30

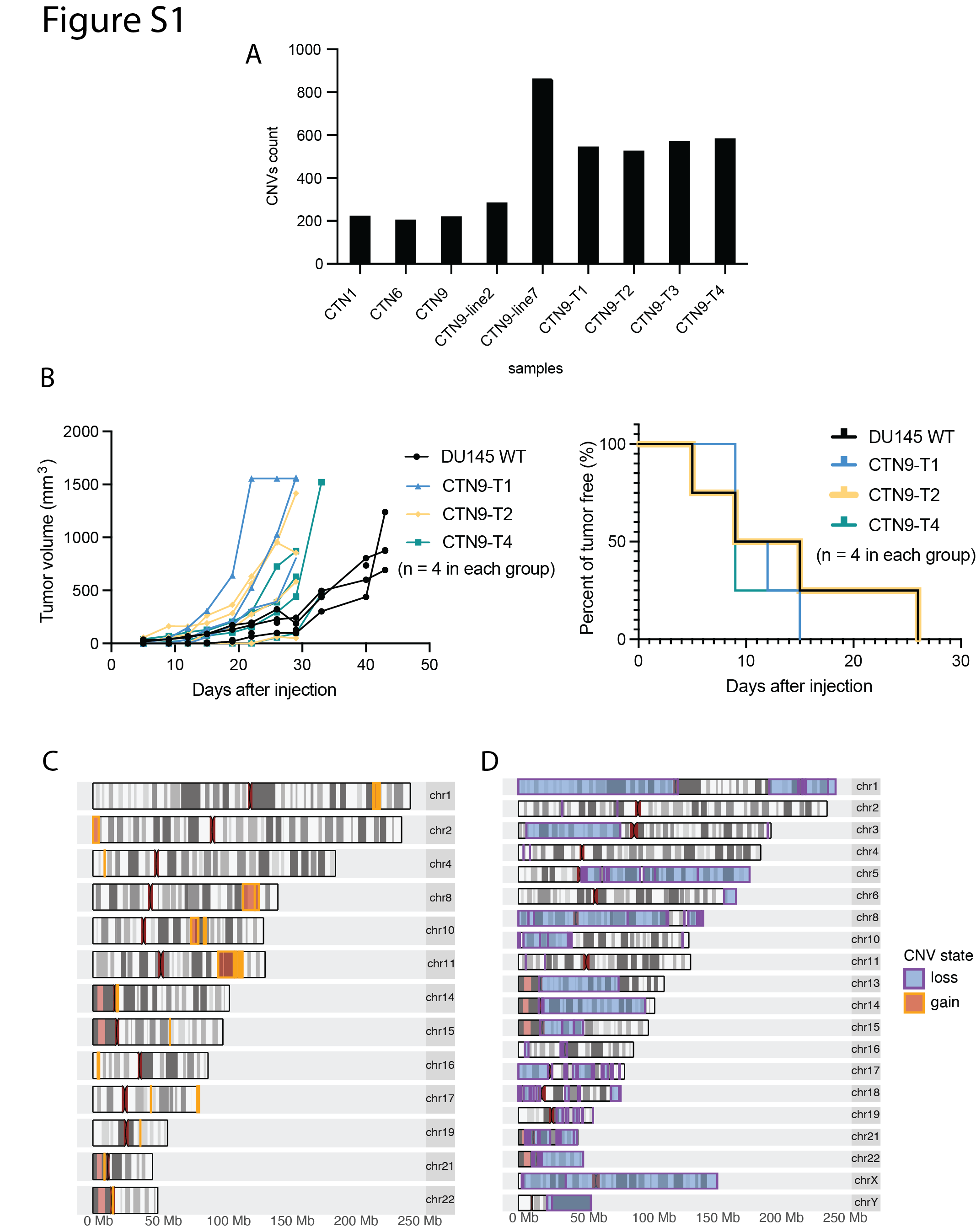

**Figure S1. CNV profiles of transient centrosome loss cells and tumor lines**

(A) Graph shows total CNV counts for the indicated cell lines.

(B) Graphs show tumor volume (left panel) and percent tumor free growth (right panel) of the CTN9 tumors lines (T1, 2, 4 and DU145, a metastatic prostate cancer cell line, as positive control) that were reinjected to form secondary xenograft tumors.

(C and D) Genomic coordinates of consistent copy number gains (C) and losses (D) across chromosomes in xenograft tumor lines CTN9-T1, CTN9-T2, CTN9-T3, and CTN9-T4. Copy number gains are highlighted in orange and losses are highlighted in blue.

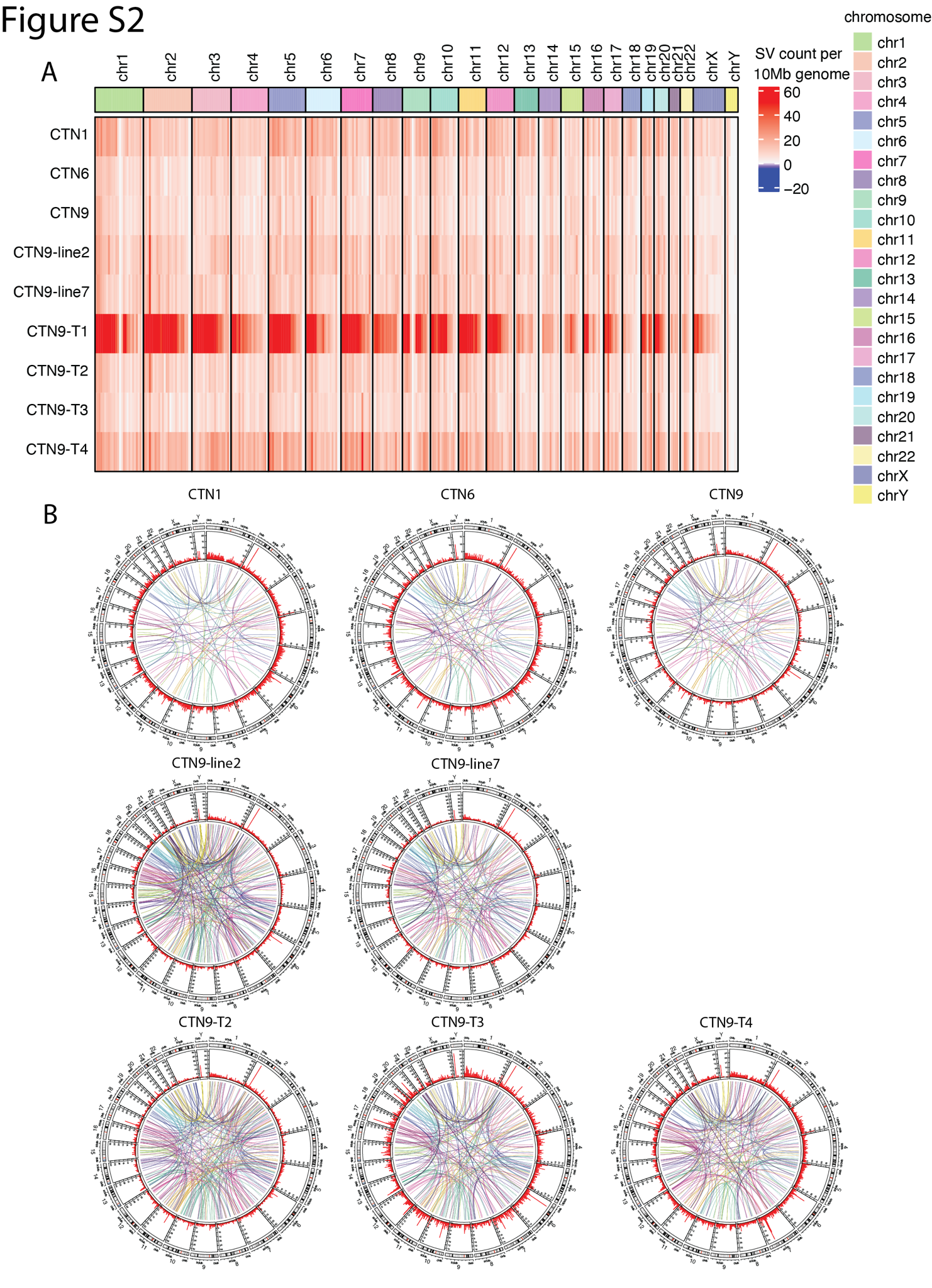

**Figure S2. Structural variation profiles of transient centrosome loss cells and tumor lines**

(A) Heatmap displaying the structural variation (SV) counts (including inversions, deletions, and duplications) across every 1 Mbp of the genome in the indicated lines. The color gradients represent the z-scored cumulative SV counts for each of the nine indicated lines.

(B) Circos plots illustrating SVs in transient centrosome loss cells (CTN1, CTN6, and CTN9), clonal lines derived from CTN9 (CTN9-line2 and CTN9-line7), and lines isolated from xenograft tumors generated from CTN9 (CTN9-T2, CTN9-T3, CTN9-T4). The innermost link plots depict translocation events, while the outer layer bar plots show the counts of SVs across every 1 Mbp of the genome.

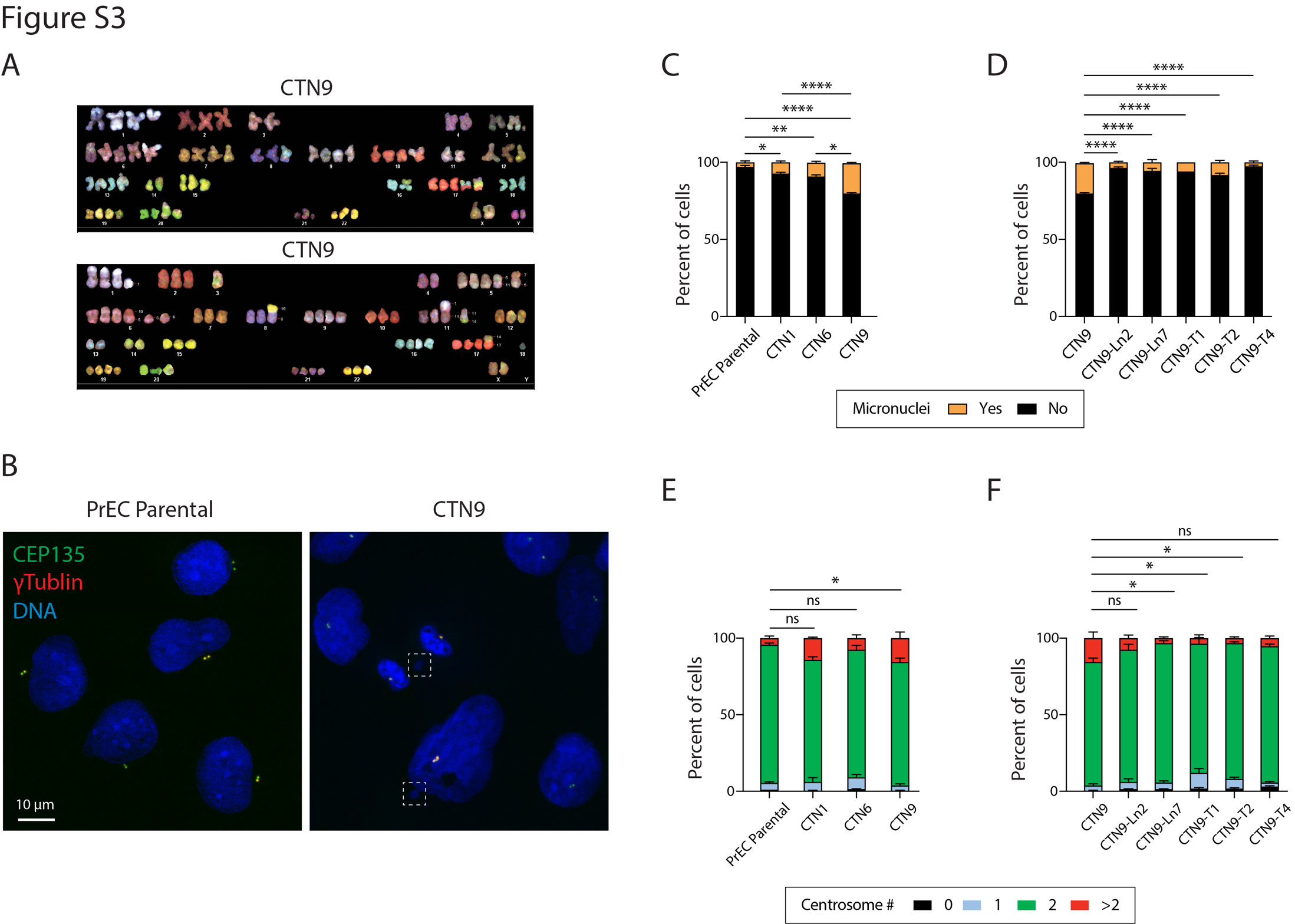

**Figure S3. CTN9 cells continue to experience CIN even after restoring centrosome numbers**

(A) Representative Spectral Karyotype (SKY) analysis of mitotic CTN9 cells.

(B) Images show PrEC and CTN9 cells stained for centrosomes. The colocalization of CEP135 (green) and γTubulin (red) mark bona fide centrosomes. DNA, blue. Dashed boxes denote micronuclei in CTN9 cells.

(C and D) Graphs show the frequency of micronuclei in the indicated lines. Data are means ± SEM. *n* = 100 cells in each of 3 experiments; significance determined using One-way ANOVA followed by Tukey’s post-hoc test for pairwise comparisons. *, *P* < 0.05; **, *P* < 0.01; ***, *P* < 0.001; ****, *P* < 0.0001; ns, not significant.

(E and F) Graphs show the average number of centrosomes (detected as co-localization of CEP135 foci and γ-Tubulin foci) per cell in the indicated cell lines. Cells with >2 centrosomes indicate centrosome amplification. Data are means ± SEM. *n* = 100 cells in each of 3 experiments; significance determined using One-way ANOVA followed by Tukey’s post-hoc test for pairwise comparisons. *, *P* < 0.05; **, *P* < 0.01; ***, *P* < 0.001; ****, *P* < 0.0001; ns, not significant.

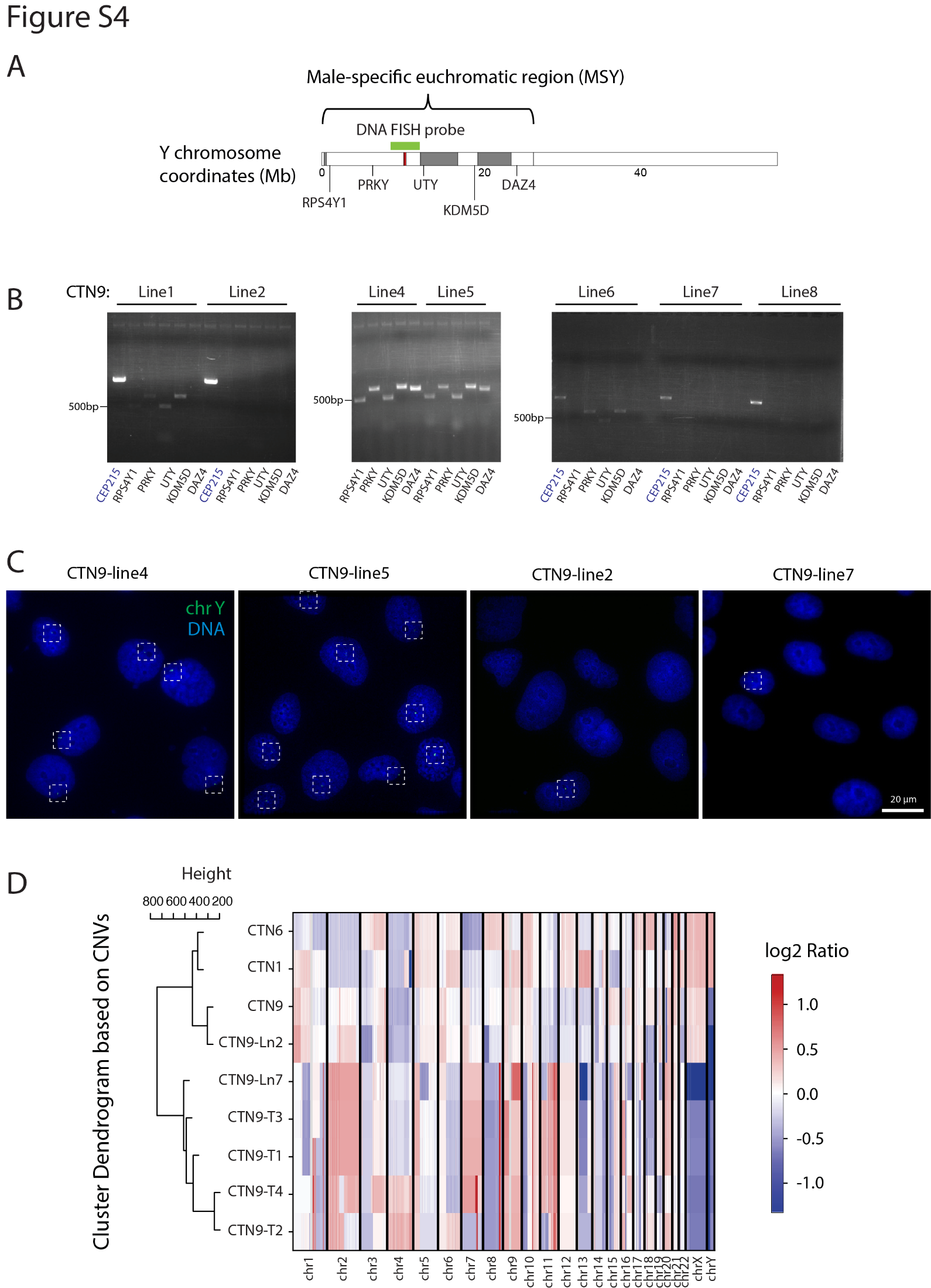

**Figure S4. Genomic heterogeneity within the CTN9 line**

(A) Schematic of Y chromosome coordinates with annotated regions targeted by PCR and FISH probes.

(B) PCR products from seven clonal clones expanded from CTN9 cells. The five gene targets (*RPS4Y1, PRKY, UTY, KDM5D, and DAZ4*) span the full length of MSY region.

(C) Fluorescent In Situ Hybridization (FISH) using a Y chromosome-specific probe (green) in cell lines derived from CTN9. Dashed boxes highlight signal in nuclei.

(D) Somatic copy number variation (CNV) heatmap of CTN9-lines 2 and 7 compared to the other lines examined in this study. The *x*-axis represents genomic region across chromosomes, and the *y*-axis indicates log2-transformed absolute copy number alteration compared to parental cells. Copy number gains (red) and losses (blue) are indicated. Hierarchical clustering dendrogram of CTN samples based on autosomal and chrX CNVs.

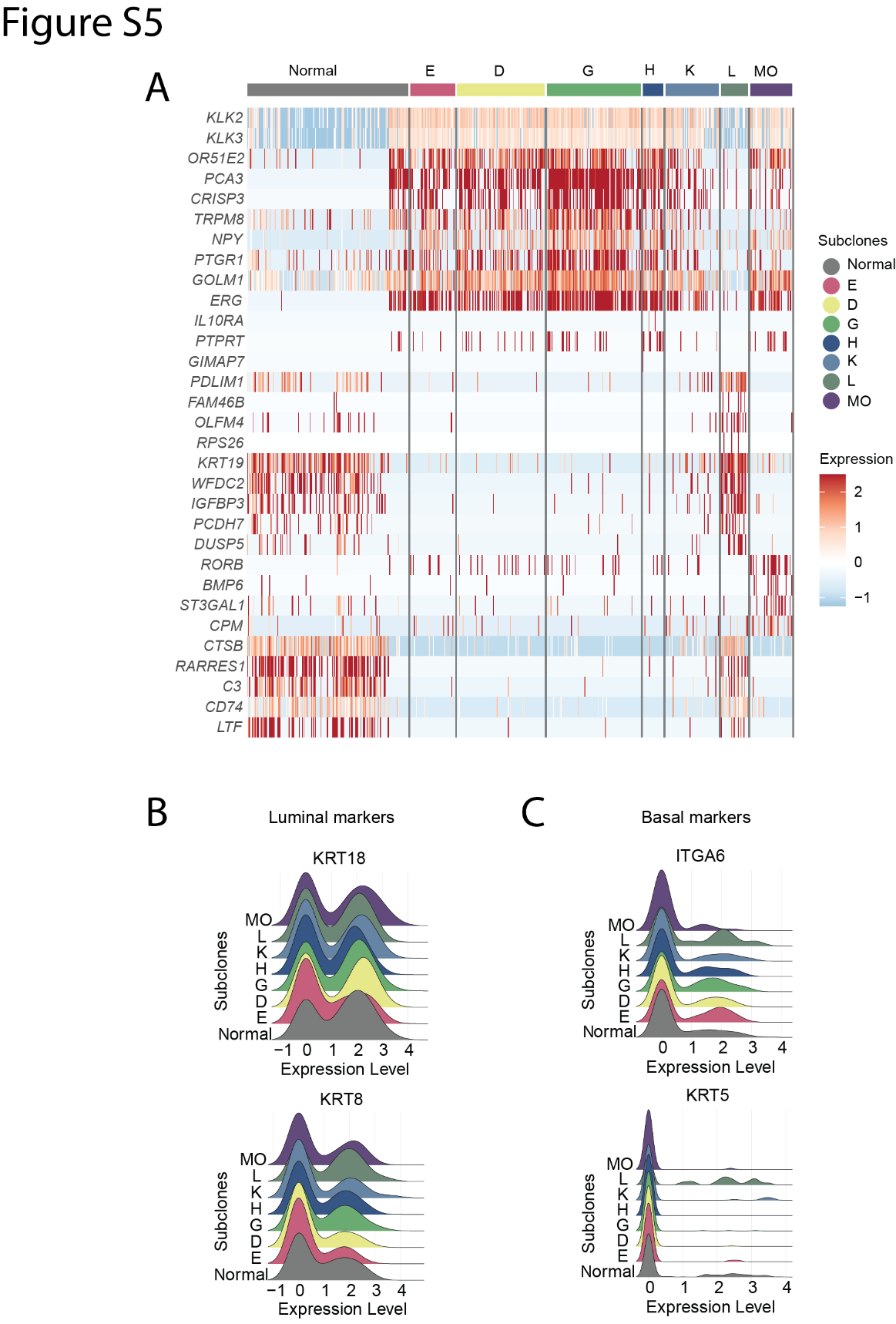

**Figure S5. Differentially expressed genes in subclones of PCa patient sample PR5249.**

(A) Transcriptomic analysis was performed on subclones, and the FindAllMakers function was used to identify gene markers with an *P*_adj_ ≤ 0.05. Only the top 10 markers for each subclone were selected for visualization. The heatmap color scale reflects normalized gene expression levels, with high expression shown in red and low expression in blue. Column annotations indicate the respective subclones.

(B and C) Ridge plots of epithelial cells from PR5249 sample, displaying the expression levels of (B) luminal marker genes and (C) basal marker genes.

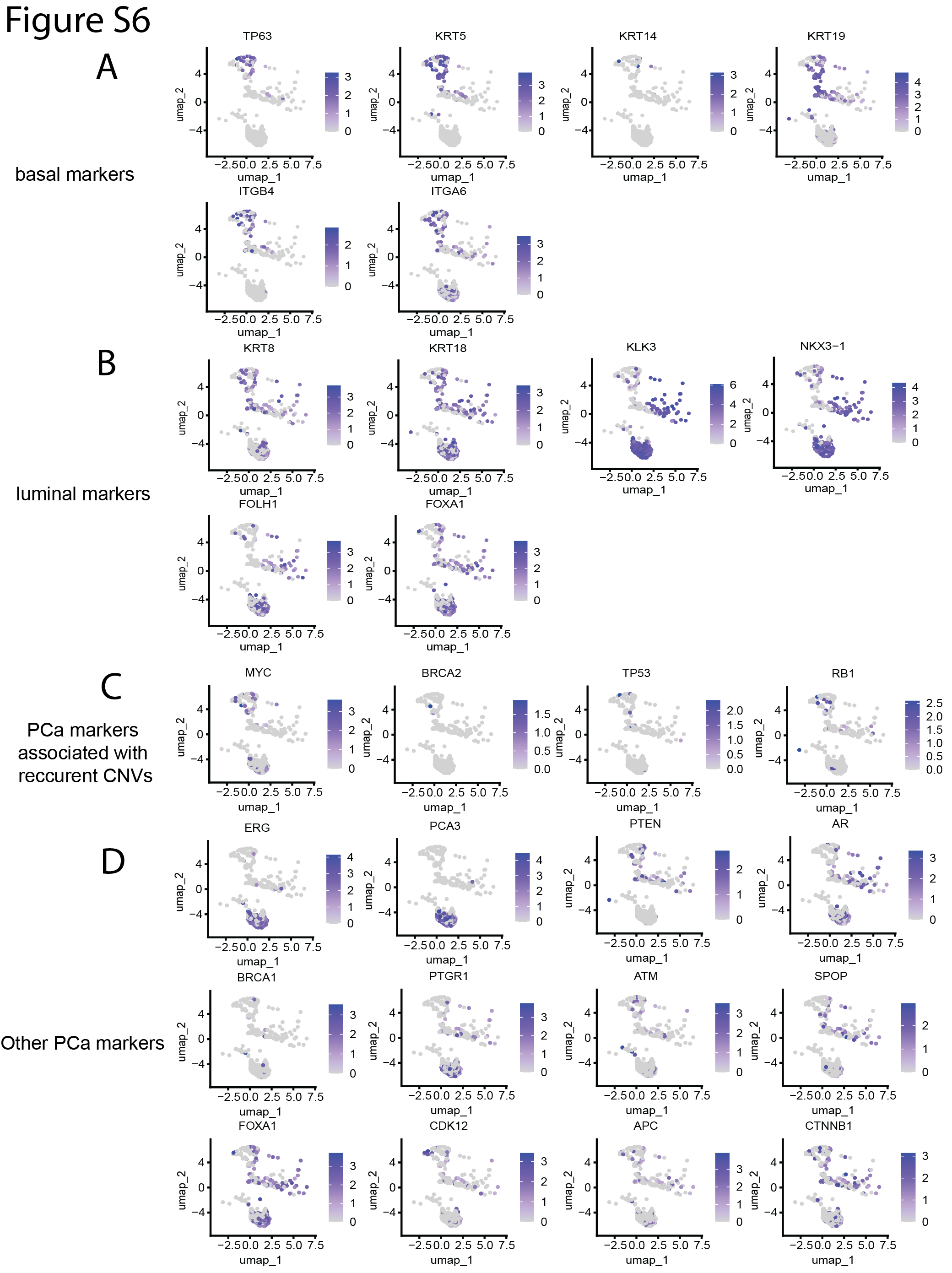

**Figure S6. UMAP visualization of single-cell data highlighting expression of PR5249 epithelial cell and PCa markers**

(A-D) UMAP was used to reduce the dimensionality of single-cell transcriptomic data, with each point representing an individual cell. Gene expression levels for specific markers are overlaid on the UMAP plot, with color intensity reflecting normalized expression (low expression in light gray, high expression in dark purple). Key gene markers under each category are labeled to illustrate their spatial distribution across cell subclones.

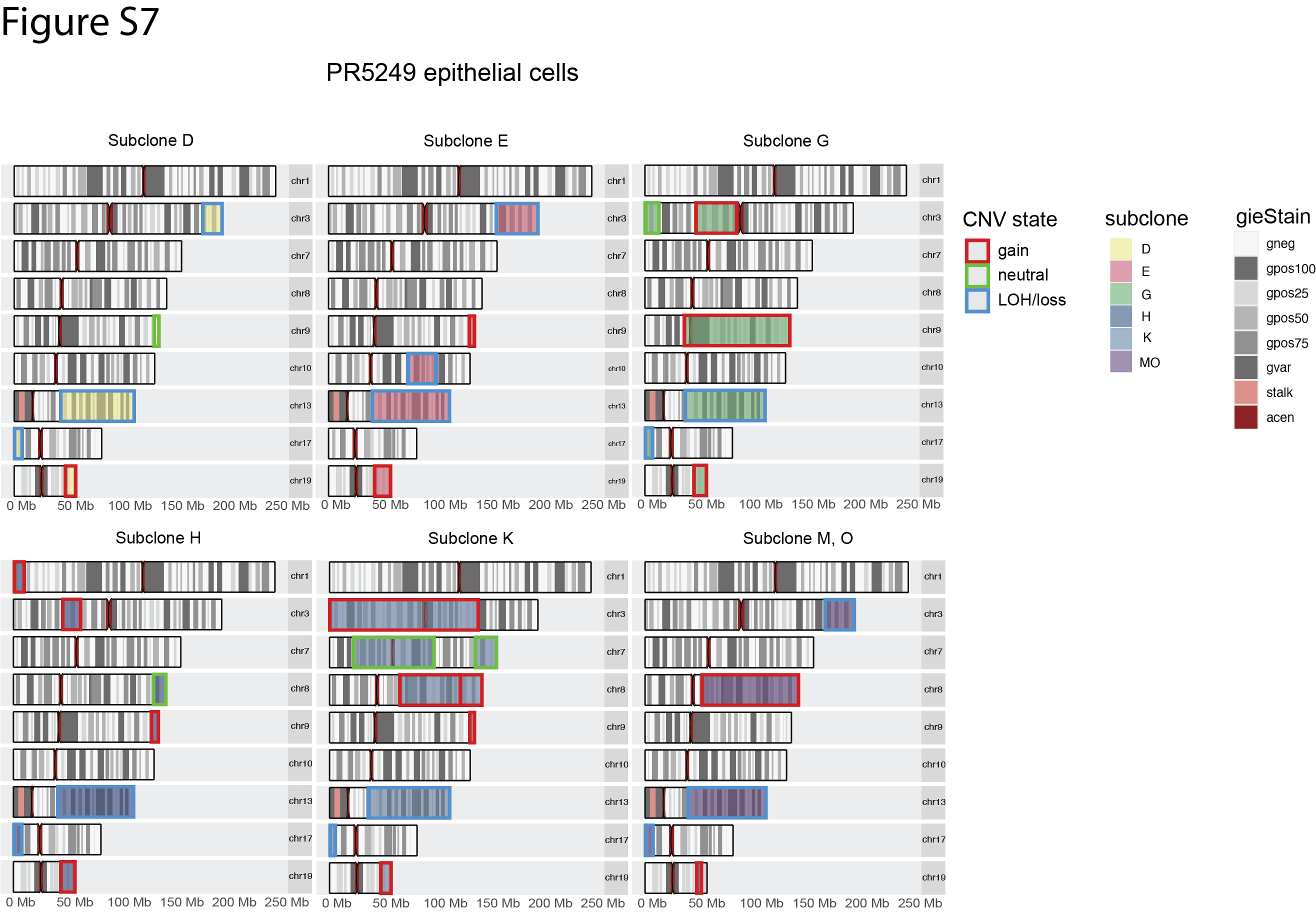

**Figure S7. Regions of CNV within distinct subclones of PCa patient sample PR5249.**

Cytogenic maps of consensus CNV regions identified in epithelia cell colonies derived from the PR5249 tumor sample, with line color indicate for CNV gain (red), neutral (green) and LOH/loss (blue) from InferCNV prediction.

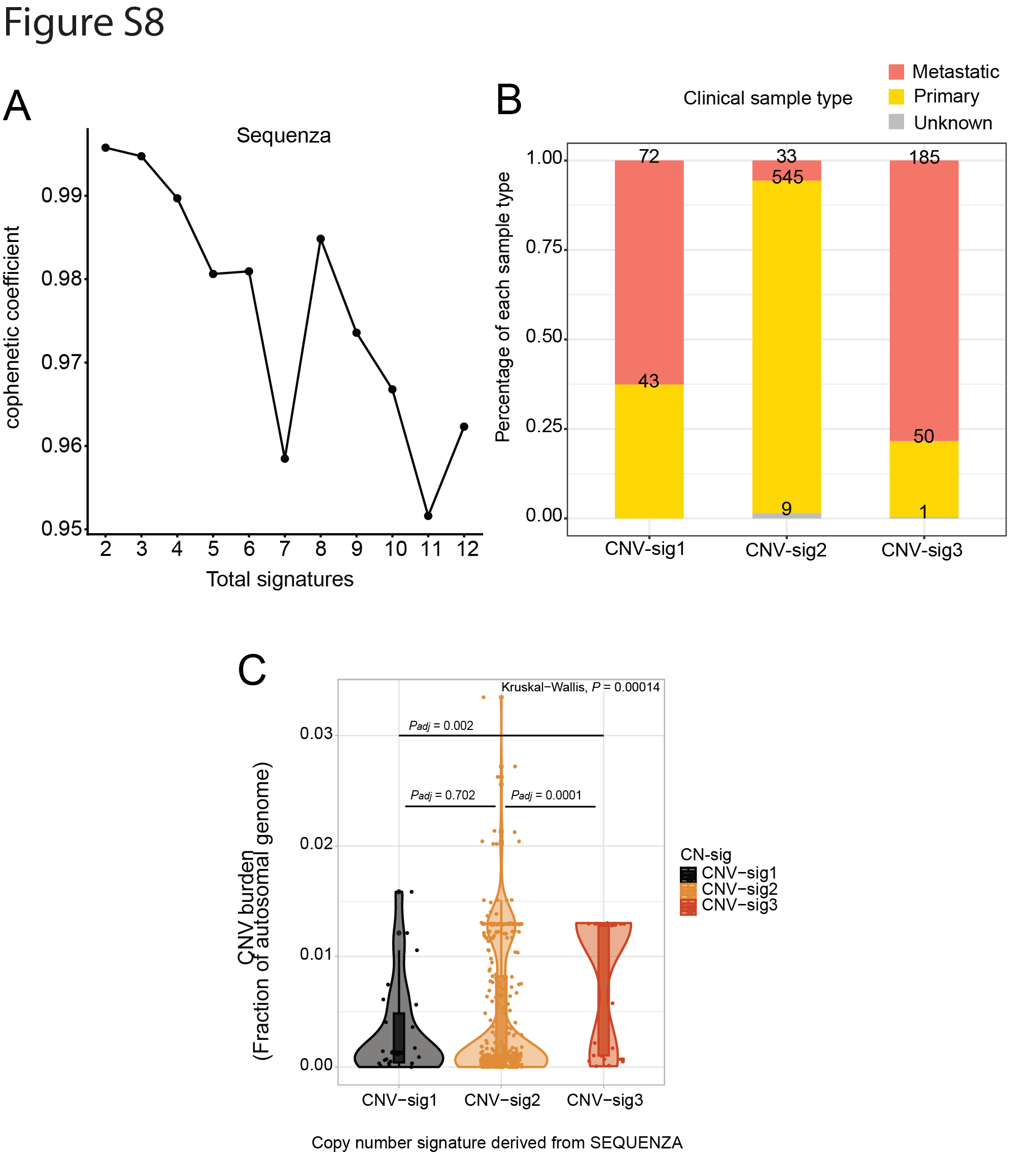

**Figure S8.** **CNV signature discovering Method validation**

(A) Elbow plot shows the cophenetic coefficient analysis for determining the optimal number of signatures, evaluated across 2 to 12 potential signatures using an input matrix derived from CNV data generated by Sequenza.

(B) Distribution of clinical sample types (metastatic vs. primary) within each copy number signature. Three CNV-signatures were identified using CNVs derived from Sequenza. The proportion of metastatic and primary samples within each signature is represented by bar plots with sample counts annotated above each bar. CNV-signature 3 exhibits the highest ratio of metastatic samples.

(C) Violin plot illustrating the CNV burden across the three CNV-signatures. Data were analyzed using one-way ANOVA revealing a significant overall difference among groups (*P* = 0.00014).

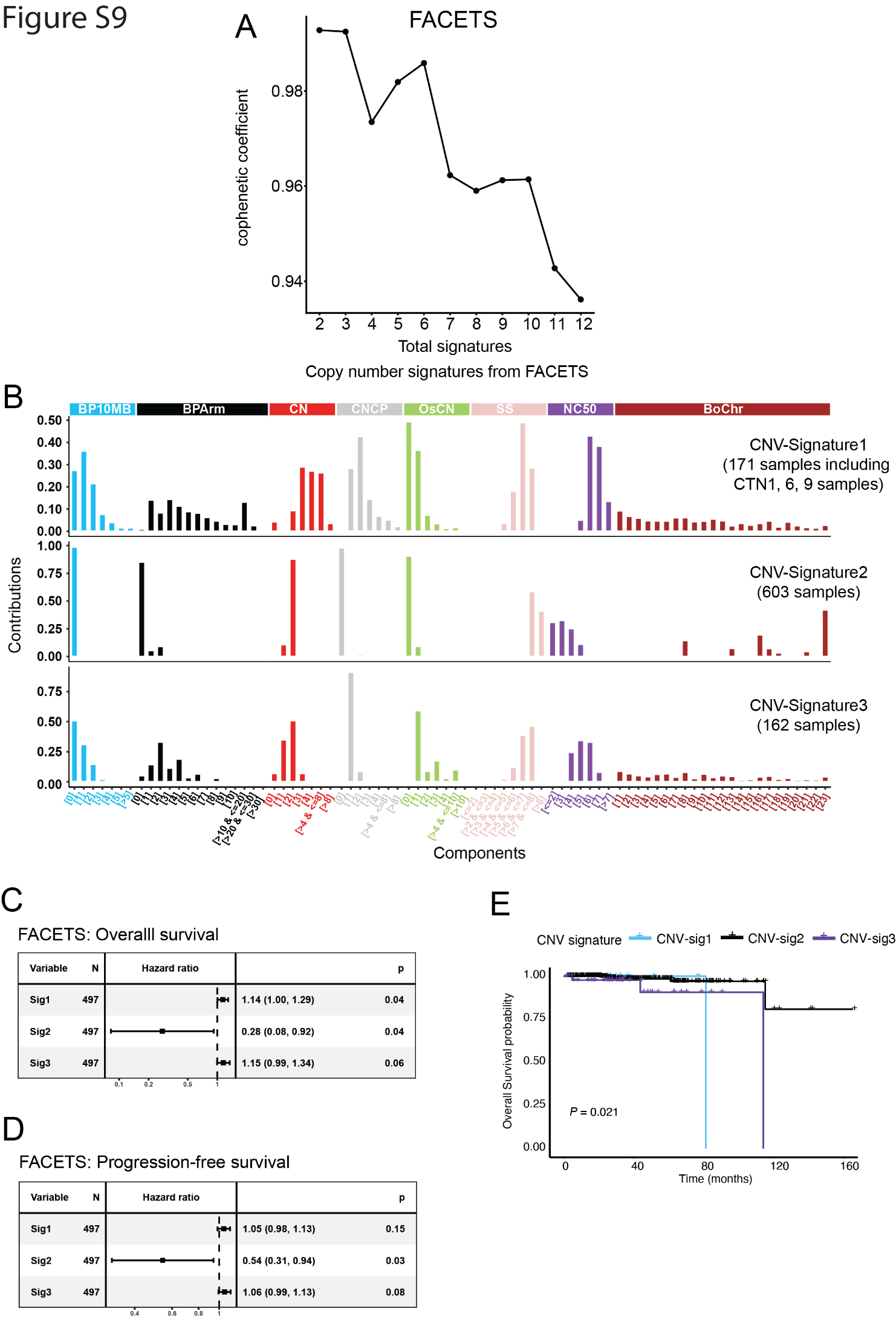

**Figure S9. CNV in transient centrosome loss cells associate with a CNV signature in PCa patients using FACETS algorithm**

(A) Elbow plot shows cophenetic coefficient analysis to determine the optimal number of signatures. The analysis was performed across 2 to 12 potential signatures using an input matrix derived from CNV data generated by FACETS.

(B) Three CNV signatures were identified from PRAD and transient centrosome-loss cell data. Each CNV signature comprises 8 distinct genomic features with a total of 80 components and were row normalized within each feature.

(C and D) Forest plots show the relative risk of CNV signature exposure for (C) overall survival and (D) progression-free survival (PFS) in 497 TCGA PRAD tumor samples. Exposure for CNV signatures was normalized to a range of 1-20 to evaluate the hazard ratio per 5% exposure increase. Hazard ratios and p-values were calculated using univariable Cox analysis. Squares represent hazard ratios and horizontal lines indicate the 95% confidence intervals. Note, transient centrosome loss samples were assigned to CNV-Signature 1 which associates with a higher hazard ratio of poor overall survival (HR = 1.14 [1.00, 1.29], *P* = 0.04). However, the association with PFS was not significant (HR = 1.05 [0.98, 1.13], P = 0.15).

(E) Kaplan-Meier survival curves comparing overall survival for TCGA-PRAD samples attributed to the three distinct CNV signatures. Statistical significance was determined using the log-rank test. The overall comparison showed significant differences among groups (*P* = 0.021).

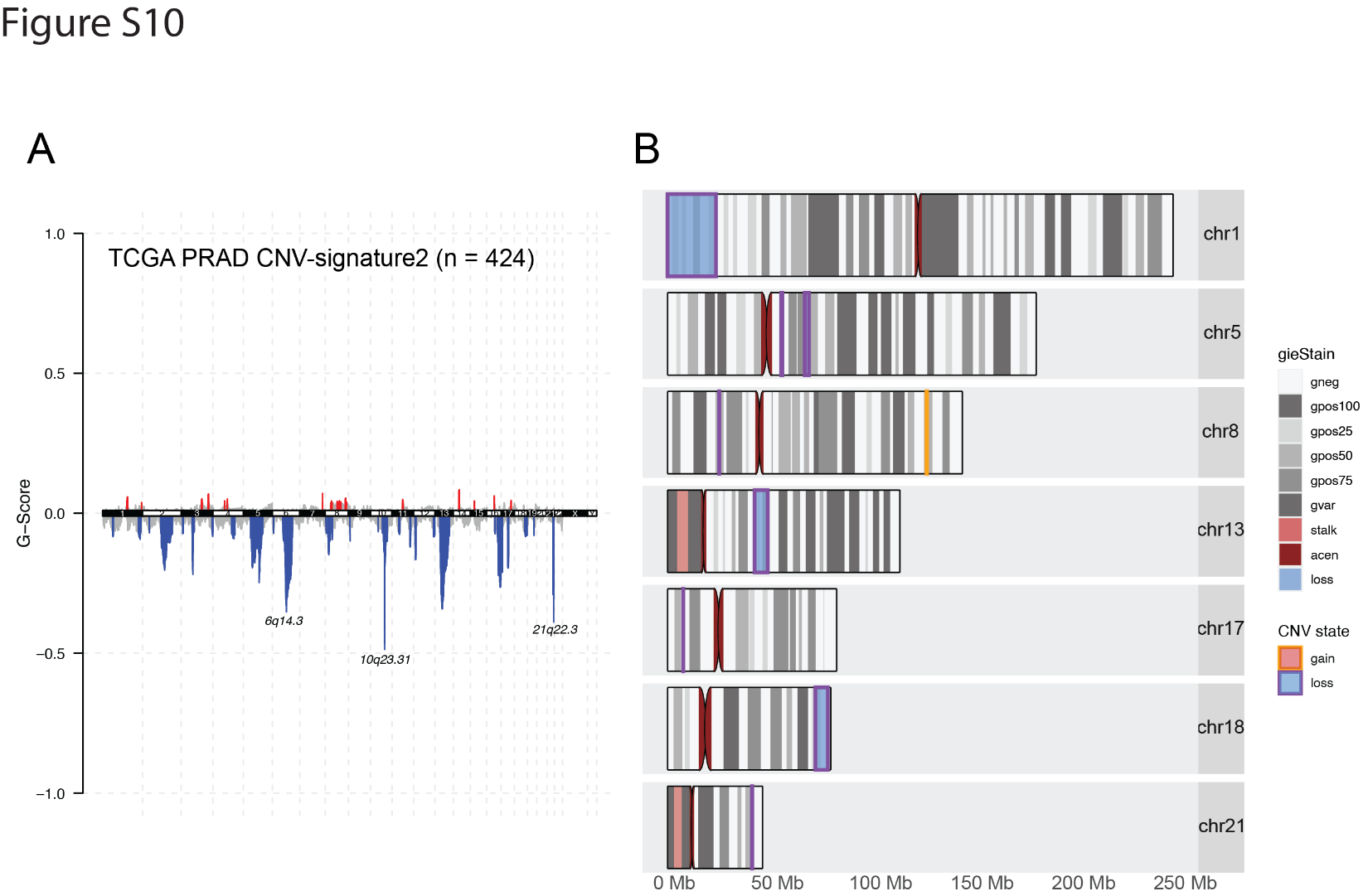

**Figure S10. Shared focal CNVs between transient centrosome loss samples and TCGA PRAD CNV-signature 3 samples**

(A) GISTIC2.0 analysis of focal CNV regions in TCGA-PRAD samples classified under CNV-signature 2 (CNV-signatures 1 and 3 are shown in Figure 4G and H).

(B) Consensus CNV regions between transient centrosome loss tumor samples (CTN9-T1-4) and TCGA PRAD CNV-signature 3-associated CNV regions (q-value < 0.1 as the cut-off for included CNV regions). Chromosomal amplifications (blue) and deletions (orange) are shown. Significant amplification and deletion regions identified by GISTIC2.0 overlapped with CNVs showing consistent directionality across all four CTN9-derived tumor samples.

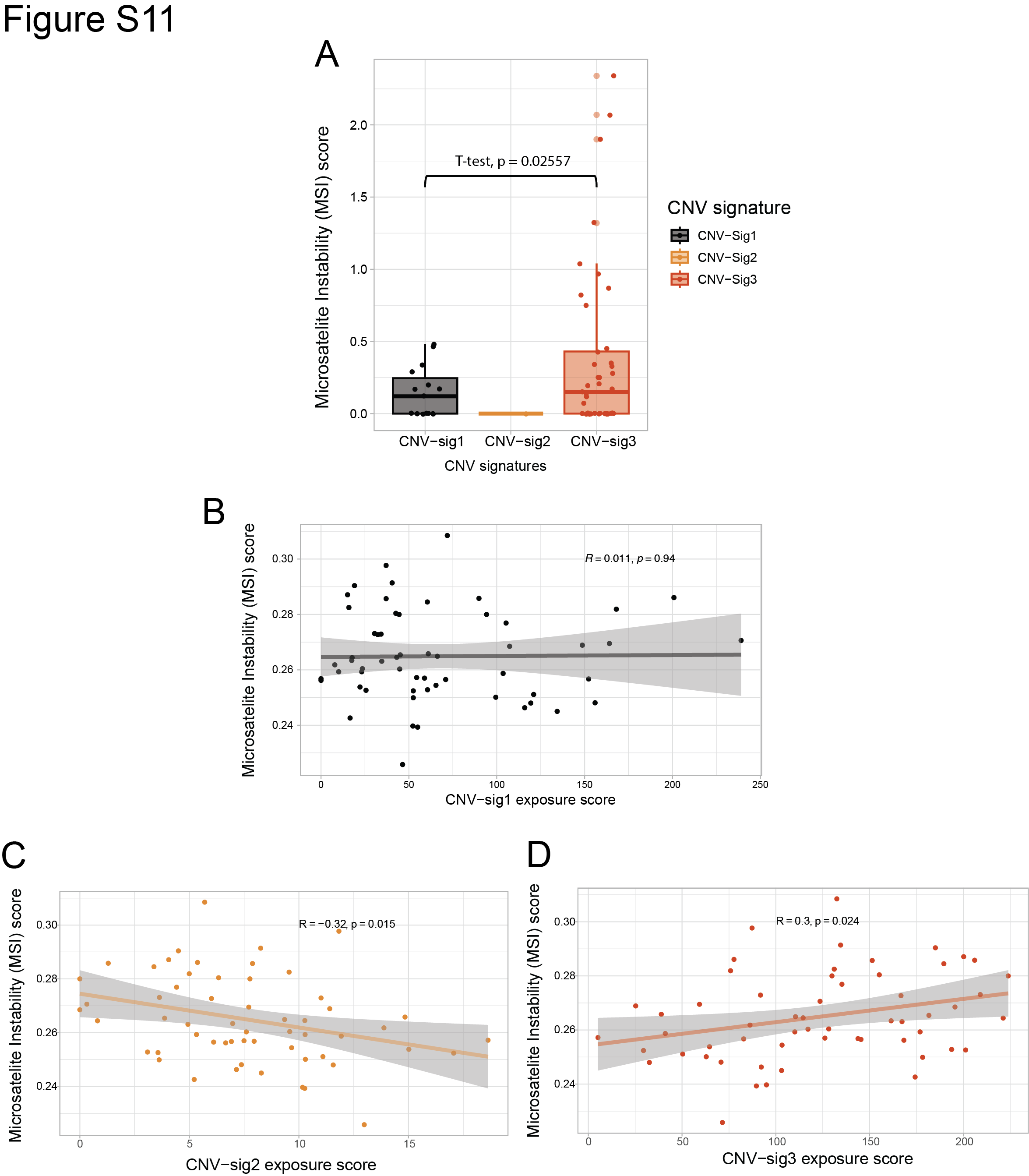

**Figure S11. Association between transient centrosome-loss associated CNV-signature 3 and microsatellite instability (MSI) score.**

(A) Box plot showing the distribution of MSI scores in SU2C/PCF PRAD samples across the three previously defined CNV-signatures. Samples with CNV-signature 3 exhibit significantly higher MSI scores compared to those with CNV-signature 1 (t-test, *P* = 0.02557). Statistical comparisons involving CNV-signature 2 were not feasible due to the presence of only one sample in this group and the Kruskal-Wallis test across all three signatures could not be applied for the same reason.

(B-D) Pearson correlation coefficient between CNV-signatures exposure scores and MSI scores, highlighted the relationship between CNV-signature activity and microsatellite instability.

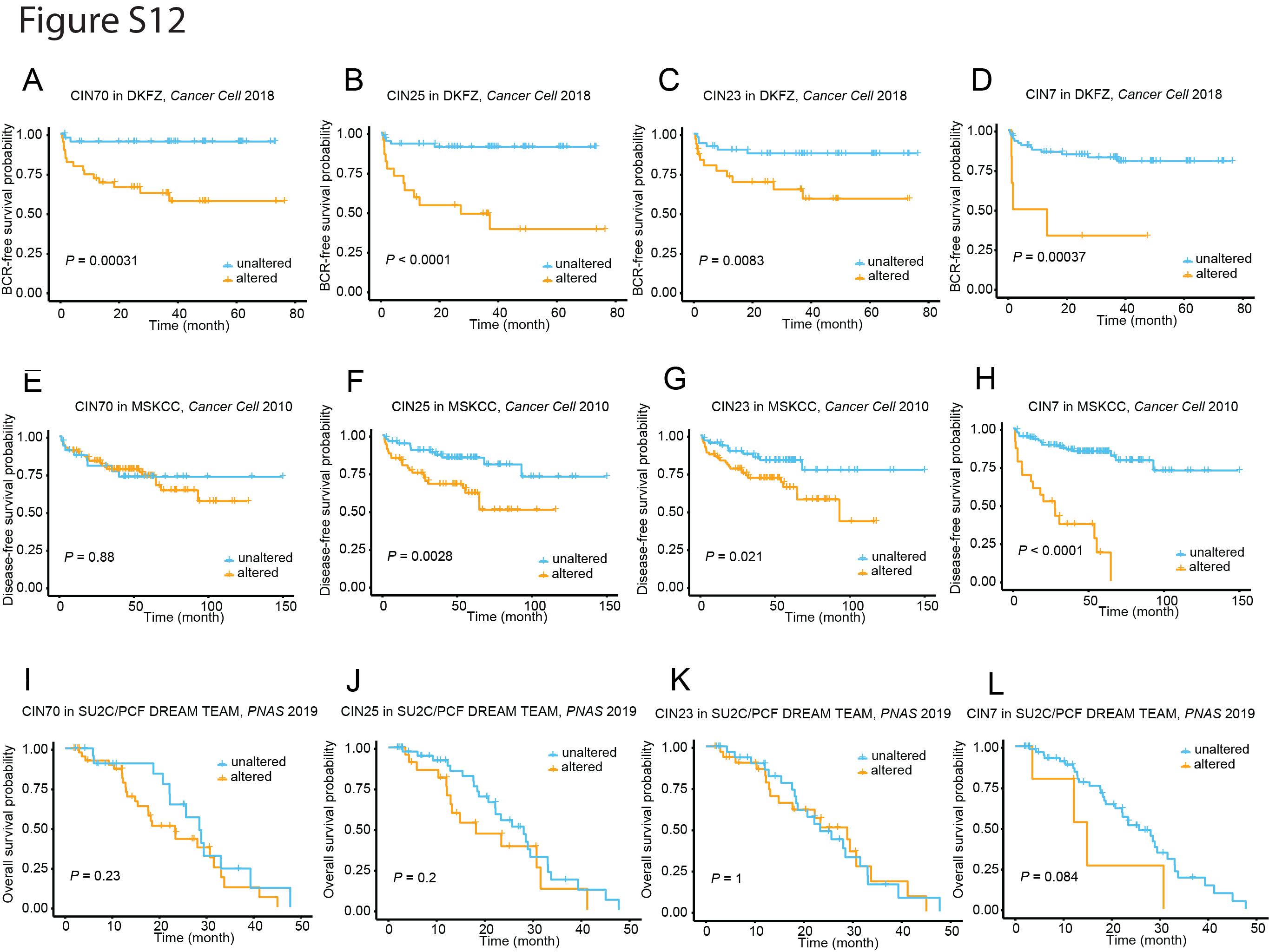

**Figure S12. A prediction of clinical outcomes using different transcriptional CIN signature gene sets.**

(A-L) Five different CIN signature gene sets were used to classify samples and predict clinical outcomes following the same methodology applied to CNV-signature 3-derived CIN9 gene expression signature. Results from the CA20 gene set are not included in this figure due to space constraints.

(A-D) Kaplan-Meier curves showing biomedical recurrence-free survival in the primary PCa DFKZ cohort from Gerhauser et al.^1^  Patients with dysregulated expression levels of individual CIN gene set were compared to the remaining cohort. Significant differences in biochemical recurrence-free survival were observed for all gene sets. (E - H) Results in the PCa MSK cohort from Taylor B et al.^2^

(I - L) Results in the PCa SU2C/PCF DREAN TEAM cohort from Abida W et al.^3^

**Table S1**

**Transient centrosome loss sample purity**

| **Sample** | **Tumor purity** |
| --- | --- |
| CTN-1 | 0.55 |
| CTN-6 | 0.54 |
| CTN-9 | 0.45 |
| CTN9-T1 | 0.91 |
| CTN9-T2 | 0.91 |
| CTN9-T3 | 0.91 |
| CTN9-T4 | 0.90 |

**Table S2**

**PR5249 tumor sample epithelial cells subclone CNVs and population size**

| **Subclone** | **Population size**  **(% of all events)** |
| --- | --- |
| D | 22 |
| E | 11 |
| G | 24 |
| H | 5 |
| K | 13 |
| MO | 10 |

**Table S3**

**Top 15 CNV regions enriched in CNV-signature 3 TCGA PRAD samples.**

| **CNV state** | **band** | **start** | **end** | **Involved genes** | **q-value** |
| --- | --- | --- | --- | --- | --- |
| Deletion | 13q14.12 | 42850052 | 49224037 | *EPSTI1/DNAJC15/ENOX1/CCDC122/LACC1/SMIM2/SERP2/TSC22D1/NUFIP1/GPALPP1/GTF2F2/KCTD4/TPT1/SLC25A30/COG3/ERICH6B/CBY2/SIAH3/ZC3H13/CPB2/LCP1/LRRC63/RUBCNL/LRCH1/ESD/HTR2A/SUCLA2/NUDT15/MED4/ITM2B/RB1/LPAR6/RCBTB2/CYSLTR2/FNDC3A/MLNR* | 1.6041e-06 |
| Deletion | 21q22.3 | 41455552 | 41680542 | *MX1/TMPRSS2* | 1.7041e-06 |
| Deletion | 10q23.31 | 87857401 | 88589061 | *KLLN/PTEN/RNLS/LIPJ* | 4.4389e-05 |
| Deletion | 5q11.2 | 55922849 | 56458287 | *IL31RA/IL6ST/ANKRD55`* | 0.00032273 |
| Deletion | 8p21.2 | 25403324 | 25430175 | *DOCK5/GNRH1/KCTD9* | 0.0019462 |
| Amplification | 11q13.3 | 68818519 | 69808837 | *CPT1A/MRPL21/IGHMBP2/MRGPRD/MRGPRF/TPCN2/SMIM38/MYEOV/CCND1/LTO1/FGF19/FGF4* | 0.0026131 |
| Deletion | 6q15 | 79703209 | 116250412 | *SH3BGRL2/ELOVL4/TTK/BCKDHB/TENT5A/IBTK/TPBG/UBE3D/DOP1A/PGM3/RWDD2A/ME1/PRSS35/SNAP91/RIPPLY2/CYB5R4/MRAP2/CEP162/TBX18/NT5E/SNX14/SYNCRIP/HTR1E/CGA/ZNF292/GJB7/SMIM8/C6orf163/CFAP206//SLC35A1/RARS2/ORC3/AKIRIN2/SPACA1/CNR1/RNGTT/PNRC1/SRSF12/PM20D2/GABRR1/GABRR2/UBE2J1/RRAGD/ANKRD6/LYRM2/MDN1/CASP8AP2/GJA10/BACH2/MAP3K7/EPHA7/MANEA/FUT9/UFL1/FHL5/GPR63/NDUFAF4/KLHL32/MMS22L/POU3F2/FBXL4/FAXC/COQ3/PNISR/USP45/TSTD3/CCNC/PRDM13/MCHR2/SIM1/ASCC3/GRIK2/HACE1/LIN28B/BVES/POPDC3/PREP/PRDM1/ATG5/CRYBG1/RTN4IP1/QRSL1/CD24/MTRES1/BEND3/PDSS2/SOBP/SCML4/SEC63/OSTM1/NR2E1/SNX3/AFG1L/FOXO3/ARMC2/SESN1/CEP57L1/CD164/PPIL6/SMPD2/MICAL1/ZBTB24/AK9/FIG4/GPR6/WASF1/CDC40/METTL24/DDO/SLC22A16/CDK19/AMD1/GTF3C6/RPF2/SLC16A10/MFSD4B/REV3L/TRAF3IP2/FYN/CCN6/TUBE1/FAM229B/LAMA4//RFPL4B/MARCKS/HDAC2/HS3ST5/FRK/NT5DC1/COL10A1/TSPYL4* | 0.004504 |
| Deletion | 16q24.1 | 75655647 | 90338345 | *TERF2IP/CPHXL2/DUXB/CPHXL/CNTNAP4/MON1B/SYCE1L/ADAMTS18/NUDT7/VAT1L/CLEC3A/WWOX/MAF/DYNLRB2/CDYL2/CMC2/CENPN/ATMIN/C16orf46/GCSH/PKD1L2/BCO1/GAN/CMIP/PLCG2/SDR42E1/HSD17B2/MPHOSPH6/CDH13/HSBP1/MLYCD/OSGIN1/NECAB2/SLC38A8/MBTPS1/HSDL1/DNAAF1/TAF1C/ADAD2/KCNG4/WFDC1/ATP2C2/MEAK7/COTL1/KLHL36/USP10/CRISPLD2/ZDHHC7/KIAA0513/CIBAR2/GSE1/GINS2/C16orf74/EMC8/COX4I1/IRF8/FOXF1/MTHFSD/FOXC2/FOXL1/C16orf95/FBXO31/MAP1LC3B/ZCCHC14/JPH3/KLHDC4/SLC7A5/CA5A/BANP/ZNF469/ZFPM1/ZC3H18/IL17C/CYBA/MVD/SNAI3/RNF166/CTU2/PIEZO1/CDT1/APRT/GALNS/TRAPPC2L/PABPN1L/CBFA2T3/ACSF3/CDH15/SLC22A31/ZNF778/ANKRD11/SPG7/RPL13/CPNE7/DPEP1/CHMP1A/SPATA33/CDK10/SPATA2L/VPS9D1/ZNF276/FANCA/SPIRE2/TCF25/MC1R//TUBB3/DEF8/DBNDD1/GAS8/PRDM7* | 0.004504 |
| Amplification | 8q22.1 | 97844405 | 97888659 | *LAPTM4B/MATN2* | 0.0085572 |
| Deletion | 16q22.2 | 68445754 | 74437521 | *SMPD3/ZFP90/CDH3/CDH1/TANGO6/HAS3/CHTF8/DERPC/UTP4/SNTB2//VPS4A/COG8/PDF/NIP7/TMED6/TERF2/CYB5B/NFAT5/NQO1/NOB1/WWP2/CLEC18A/PDPR/CLEC18C/EXOSC6/AARS1/DDX19B//DDX19A/ST3GAL2/FCSK/COG4/SF3B3/IL34/MTSS2/VAC14/HYDIN/CMTR2/CALB2/TLE7/ZNF23/ZNF19/CHST4/TAT/MARVELD3/PHLPP2/AP1G1/ATXN1L/IST1/ZNF821/PKD1L3/DHODH/TXNL4B/HP/HPR/DHX38/PMFBP1/ZFHX3/PSMD7/NPIPB15/CLEC18B* | 0.010817 |
| Deletion | 12p13.1 | 8097172 | 18264383 | *NECAP1/CLEC4A/ZNF705A/FAM90A1/CLEC6A/CLEC4D/CLEC4E/AICDA/MFAP5/RIMKLB/A2ML1/PHC1/M6PR/KLRG1/A2M/PZP/KLRB1/CLEC2D/CD69/KLRF1/CLEC2B/KLRF2/CLEC2A/CLEC12A/CLEC1B/CLEC12B/CLEC9A/CLEC1A/CLEC7A/OLR1/TMEM52B/GABARAPL1/KLRD1/KLRK1/KLRC4-KLRK1/KLRC4/KLRC3/KLRC2/KLRC1/EIF2S3B/MAGOHB/STYK1/YBX3/TAS2R7/TAS2R8/TAS2R9/PRH1/TAS2R10/PRR4/TAS2R13/PRH2/TAS2R14/TAS2R50/TAS2R20/TAS2R19/TAS2R31/TAS2R46/TAS2R43/TAS2R30/SMIM10L1/TAS2R42/PRB3/PRB4/PRB1/PRB2/ETV6/BCL2L14/LRP6/MANSC1/BORCS5/DUSP16/CREBL2/GPR19/CDKN1B/APOLD1/DDX47/GPRC5A/GPRC5D/HEBP1/FAM234B/GSG1/EMP1/GRIN2B/ATF7IP/PLBD1/GUCY2C/H4C16/H2AJ/WBP11/C12orf60/SMCO3/ART4/MGP/ERP27/ARHGDIB/PDE6H/RERG/PTPRO/EPS8/STRAP/DERA/SLC15A5/MGST1/LMO3/RERGL/PIK3C2G* | 0.017143 |
| Deletion | 5q13.2 | 67196585 | 73454225 | *CD180/PIK3R1/SLC30A5/CCNB1/CENPH/MRPS36/CDK7/CCDC125/AK6/TAF9/RAD17/MARVELD2/OCLN/GTF2H2C/SERF1B/SMN2/SERF1A/SMN1/NAIP/GTF2H2/BDP1/MCCC2/CARTPT/MAP1B/MRPS27/PTCD2/ZNF366/TNPO1/FCHO2/TMEM171/TMEM174/FOXD1* | 0.019102 |
| Amplification | 8q21.13 | 80608003 | 81797932 | *ZNF704/PAG1/FABP5/PMP2/FABP9/FABP4/FABP12/IMPA1/SLC10A5/ZFAND1/CHMP4C* | 0.0194 |
| Amplification | 8q24.21 | 127454570 | 127806762 | *MYC* | 0.039182 |
| Deletion | 17p13.1 | 7601910 | 7710389 | *FXR2/SHBG/SAT2/ATP1B2/TP53/WRAP53/EFNB3* | 0.0649 |

**Table S4**

**Consensus CNV regions enriched in CNV-signature 3 TCGA PRAD samples and centrosome loss tumor samples**

| **chromosome** | **start** | **end** | **cytobands** | **CNV state** | **Involved genes** |
| --- | --- | --- | --- | --- | --- |
| chr8 | 127454570 | 127806762 | 8q24.21 | gain | *MYC* |
| chr13 | 42850052 | 49224037 | 13q14.12 | loss | *EPSTI1/DNAJC15/ENOX1/CCDC122/LACC1/SMIM2/SERP2/TSC22D1/NUFIP1/GPALPP1/GTF2F2/KCTD4/TPT1/SLC25A30/COG3/ERICH6B/CBY2/SIAH3/ZC3H13/CPB2/LCP1/LRRC63/RUBCNL/LRCH1/ESD/HTR2A/SUCLA2/NUDT15/MED4/ITM2B/RB1/LPAR6/RCBTB2/CYSLTR2/FNDC3A/MLNR* |
| chr21 | 41455552 | 41680542 | 21q22.3 | loss | *MX1/TMPRSS2* |
| chr5 | 55922849 | 56458287 | 5q11.2 | loss | *IL31RA/IL6ST/ANKRD55* |
| chr8 | 25403324 | 25430175 | 8p21.2 | loss | *DOCK5/GNRH1/KCTD9* |
| chr17 | 7601910 | 7710389 | 17p13.1 | loss | *FXR2/SHBG/SAT2/ATP1B2/TP53/WRAP53/EFNB3* |
| chr5 | 67463165 | 69630665 | 5q13.2 | loss | *PIK3R1/SLC30A5/CCNB1/CENPH/MRPS36/CDK7/CCDC125/AK6/TAF9/RAD17/MARVELD2/OCLN/GTF2H2C* |
| chr1 | 10000 | 23783467 | 1p36.22 | loss | *OR4F5/OR4F29/OR4F16/SAMD11/NOC2L/KLHL17/PLEKHN1/PERM1/HES4/ISG15/AGRN/RNF223/C1orf159/TTLL10/TNFRSF18/TNFRSF4/SDF4/B3GALT6/C1QTNF12/UBE2J2/SCNN1D/ACAP3/PUSL1/INTS11/CPTP/TAS1R3/DVL1/MXRA8/AURKAIP1/CCNL2/MRPL20/ANKRD65/TMEM88B/VWA1/ATAD3C/ATAD3B/ATAD3A/TMEM240/SSU72/FNDC10/MIB2/MMP23B/CDK11B/SLC35E2B/CDK11A/NADK/GNB1/CALML6/TMEM52/CFAP74/GABRD/PRKCZ/FAAP20/SKI/MORN1/RER1/PEX10/PLCH2/PANK4/HES5/TNFRSF14/PRXL2B/MMEL1/TTC34/ACTRT2/PRDM16/ARHGEF16/MEGF6/TPRG1L/WRAP73/TP73/CCDC27/SMIM1/LRRC47/CEP104/DFFB/C1orf174/AJAP1/NPHP4/KCNAB2/CHD5/RPL22/RNF207/ICMT/HES3/GPR153/ACOT7/HES2/ESPN/TNFRSF25/PLEKHG5/NOL9/TAS1R1/ZBTB48/KLHL21/PHF13/THAP3/DNAJC11/CAMTA1/VAMP3/PER3/UTS2/TNFRSF9/PARK7/ERRFI1/SLC45A1/RERE/ENO1/CA6/SLC2A7/SLC2A5/GPR157/H6PD/SPSB1/SLC25A33/TMEM201/PIK3CD/CLSTN1/CTNNBIP1/LZIC/NMNAT1/RBP7/UBE4B/KIF1B/PGD/CENPS-CORT/CENPS/CORT/DFFA/PEX14/CASZ1/C1orf127/TARDBP/MASP2/SRM/EXOSC10/MTOR/ANGPTL7/UBIAD1/DISP3/FBXO2/FBXO44/MAD2L2/FBXO6/DRAXIN/AGTRAP/C1orf167/MTHFR/CLCN6/NPPA/NPPB/KIAA2013/PLOD1/MFN2/MIIP/TNFRSF8/TNFRSF1B/VPS13D/DHRS3/AADACL4/AADACL3/CFAP107/PRAMEF12/PRAMEF1/PRAMEF11/HNRNPCL1/PRAMEF2/PRAMEF4/PRAMEF10/PRAMEF7/PRAMEF6/PRAMEF27/HNRNPCL3/PRAMEF25/HNRNPCL2/PRAMEF26/HNRNPCL4/PRAMEF9/PRAMEF13/PRAMEF18/PRAMEF5/PRAMEF8/PRAMEF33/PRAMEF15/PRAMEF14/PRAMEF19/PRAMEF17/PRAMEF20/LRRC38/PDPN/PRDM2/KAZN/TMEM51/FHAD1/EFHD2/CTRC/CELA2A/CELA2B/CASP9/DNAJC16/AGMAT/DDI2/RSC1A1/PLEKHM2/SLC25A34/TMEM82/FBLIM1/UQCRHL/SPEN/ZBTB17/SRARP/HSPB7/CLCNKA/CLCNKB/FAM131C/EPHA2/ARHGEF19/CPLANE2/FBXO42/SZRD1/SPATA21/NECAP2/NBPF1/CROCC/MFAP2/ATP13A2/SDHB/PADI2/PADI1/PADI3/PADI4/PADI6/RCC2/ARHGEF10L/ACTL8/IGSF21/KLHDC7A/PAX7/TAS1R2/ALDH4A1/IFFO2/UBR4/EMC1/MRTO4/AKR7L/AKR7A3/AKR7A2/SLC66A1/CAPZB/MICOS10/NBL1/MICOS10-NBL1/HTR6/TMCO4/RNF186/OTUD3/PLA2G2E/PLA2G2A/PLA2G5/PLA2G2D/PLA2G2F/PLA2G2C/UBXN10/VWA5B1/CAMK2N1/MUL1/FAM43B/CDA/PINK1/DDOST/KIF17/SH2D5/HP1BP3/EIF4G3/ECE1/NBPF3/ALPL/RAP1GAP/USP48/LDLRAD2/HSPG2/CELA3B/CELA3A/CDC42/WNT4/ZBTB40/EPHA8/C1QA/C1QC/C1QB/EPHB2/LACTBL1/TEX46/KDM1A/LUZP1/HTR1D/HNRNPR/ZNF436/TCEA3/ASAP3/E2F2/ID3/RPL11/ELOA/PITHD1* |
| chr18 | 72734251 | 78902869 | 18q22.3 | loss | *NETO1/FBXO15/TIMM21/CYB5A/C18orf63/DIPK1C/CNDP2/CNDP1/ZNF407/PTGR3/TSHZ1/SMIM21/ZNF516/ZNF236/MBP/GALR1* |

**Table S5**

**CNV-signature 3 associated 9 CIN genes**

| **Genes** | **Chromosome** | **band** | **Start Position** | **End Position** | **Description** | **Function** |
| --- | --- | --- | --- | --- | --- | --- |
| PRR11 | 17 | q22 | 59155732 | 59206709 | proline rich 11 | cell cycle regulation |
| PPP6R3 | 11 | q13.2 | 68460731 | 68615334 | protein phosphatase 6 regulatory subunit 3 | serine and threonine phosphorylation regulation during mitosis |
| ACOT2 | 14 | q24.3 | 73567620 | 73575658 | acyl-CoA thioesterase 2 | lipid metabolism |
| SHC2 | 19 | p13.3 | 416589 | 461033 | src homology 2 domain containing transforming protein | signaling adapter |
| SMC2 | 9 | q31.1 | 104094260 | 104141419 | structural maintenance of chromosomes 2 | condensin I and II complex |
| KIF11 | 10 | q23.33 | 92574105 | 92655395 | kinesin family member 11 | mitotic spindle assembly and function |
| POLE3 | 9 | q32 | 113407235 | 113410675 | DNA polymerase epsilon 3, accessory subunit | nucleotide excision repair |
| TMEM183A | 1 | q32.1 | 203007374 | 203024848 | transmembrane protein 183A | SCF ubiquitin ligase component |
| ZNF667 | 19 | q13.43 | 56439325 | 56478065 | zinc finger protein 667 | transcription repressor |

### **Reference**

1. Gerhauser, C., Favero, F., Risch, T., Simon, R., Feuerbach, L., Assenov, Y., Heckmann, D., Sidiropoulos, N., Waszak, S.M., Hübschmann, D., et al. (2018). Molecular Evolution of Early-Onset Prostate Cancer Identifies Molecular Risk Markers and Clinical Trajectories. Cancer Cell *34*, 996-1011.e8. <https://doi.org/10.1016/j.ccell.2018.10.016>.

2. Taylor, B.S., Schultz, N., Hieronymus, H., Gopalan, A., Xiao, Y., Carver, B.S., Arora, V.K., Kaushik, P., Cerami, E., Reva, B., et al. (2010). Integrative Genomic Profiling of Human Prostate Cancer. Cancer Cell *18*, 11–22. <https://doi.org/10.1016/j.ccr.2010.05.026>.

3. Abida, W., Cyrta, J., Heller, G., Prandi, D., Armenia, J., Coleman, I., Cieslik, M., Benelli, M., Robinson, D., Allen, E.M.V., et al. (2019). Genomic correlates of clinical outcome in advanced prostate cancer. Proc. Natl. Acad. Sci. United States Am. *116*, 11428–11436. <https://doi.org/10.1073/pnas.1902651116>.
